# Cocaine-Enhanced Enkephalin Gates Plasticity of Distinct Striatal Synapses and Facilitates Cue-Induced Cocaine-Seeking During Abstinence

**DOI:** 10.64898/2026.03.11.711212

**Authors:** Kanako Matsumura, Polina Lyuboslavsky, Amelia Nicot, Bailey Remmers, Mia Smith, In Bae Choi, Gabriel Garza, Anjali Baratham, Lauren K. Dobbs

**Affiliations:** Interdisciplinary Neuroscience Program, The University of Texas at Austin, Austin, TX, USA, 78712; Department of Neuroscience, The University of Texas at Austin, Austin, TX, USA, 78712; Waggoner Center for Alcohol and Addiction Research, The University of Texas at Austin, Austin, TX, USA, 78712; Department of Neurology, Dell Medical School, The University of Texas at Austin, Austin, TX, USA, 78712

**Keywords:** striatum, enkephalin, D2-MSN, GABA, cocaine, seeking

## Abstract

**Background:** Cocaine seeking after abstinence is associated with increased ventral pallidum (VP) activity. This increase appears to involve an opioid peptide that suppresses GABA release from D2 receptor-expressing medium spiny neurons (D2-MSNs) to the VP, but its identity is unclear. We previously found that cocaine abstinence increases enkephalin mRNA expression in D2-MSNs and that exogenous enkephalin inhibits GABA release from these neurons. We therefore hypothesized that cocaine-enhanced enkephalin release from D2-MSNs suppresses GABA release into the VP, which disinhibits VP neurons and promotes cocaine seeking.

**Methods:** To examine how enkephalin alters striatal circuit plasticity and cocaine seeking following abstinence, we performed *ex vivo* electrophysiology and cocaine self-administration in mice lacking enkephalin selectively from D2-MSNs (D2-PenkKO). Recordings were obtained from the primary D2-MSNs targets: VP neurons and D1 receptor-expressing MSNs (D1-MSNs).

**Results:** Following cocaine abstinence, GABA release from D2-MSNs to VP neurons was reduced in controls, but not D2-PenkKOs. In contrast, cocaine abstinence did not alter GABA release to D1-MSNs. Optogenetically evoked GABA release from D2-MSNs inhibited VP neuron firing in saline-abstinent controls and D2-PenkKOs, but not cocaine-abstinent controls. Female, but not male, D2-PenkKOs self-administered less cocaine in early acquisition and exhibited blunted cue-induced cocaine seeking after abstinence.

**Conclusion:** Abstinence from cocaine self-administration increases D2-MSN enkephalin expression, leading to auto-inhibition of GABA release onto VP neurons, but not D1-MSNs, thereby selectively disinhibiting VP neuron activity. These findings identify enkephalin as a key mediator of cocaine abstinence-induced striatopallidal circuit plasticity and suggest that this plasticity contributes to cocaine seeking following abstinence.

## Introduction

Cocaine use disorder is a costly and prevalent public health challenge marked by high rates of relapse despite prolonged abstinence. Clinical and preclinical evidence suggests the reactivity to drug-related cues and relapse to drug seeking intensifies over drug abstinence (1–6). Reinstatement of reward seeking, including cocaine seeking, is attenuated by lesioning or inactivating the VP, indicating that VP activity is necessary for cocaine seeking (7–10). Additionally, VP neuron activity, inferred using calcium indicators, is heightened during cue-induced cocaine seeking (11). Prior work has identified the inhibition of GABA release from the striatum to the ventral pallidum (VP) as a circuit mechanism driving VP activity, because chemogenetic inhibition of dopamine D2 receptor-expressing MSNs (D2-MSNs) increases cue-induced cocaine seeking, and this enhanced seeking is absent when VP neurons are inhibited (10). Collectively, these studies suggest that reduced GABA release from D2-MSNs onto VP neurons disinhibits the VP, and that this circuit adaptation drives reinstatement of cocaine-seeking.

Evidence from *ex vivo* electrophysiology suggests striatal opioid peptides mediate suppressed GABA release from D2-MSNs. Following extinction from cocaine self-administration, spontaneous and electrically-evoked GABA release onto VP neurons is reduced compared with yoked saline controls (12). This GABA suppression is reversed by the µ-opioid receptor (MOR) antagonist CTOP, suggesting increased opioid peptide tone (12). Similarly, optogenetically-evoked GABA release from D2-MSNs is reversed by CTOP or the δ-opioid receptor (DOR) antagonist naltrindole (10,13). Together, this evidence strongly suggests that abstinence from long-term cocaine exposure chronically inhibits GABA release from D2-MSNs onto VP neurons via heightened opioid peptide tone acting on MORs or DORs in D2-MSNs.

The identity and source of the opioid peptide that regulates GABA release from D2-MSNs following cocaine abstinence is unknown. One candidate is enkephalin, which is enriched in D2-MSNs, has a high affinity for MORs and DORs (14,15), and inhibits GABA release from D2-MSNs onto neighboring MSNs when exogenously applied (16,17). Moreover, cocaine abstinence increases abundance of striatal enkephalin mRNA and peptide (17–20). However, it is unknown if striatal enkephalin is responsible for the striatopallidal circuit GABA plasticity driving enhanced cocaine seeking during abstinence. Further, it is not known if cocaine abstinence similarly suppresses GABA release from D2-MSNs onto D1 receptor-expressing MSNs (D1-MSNs). Altered GABA release in this intrastriatal circuit may promote cocaine seeking as we previously found it undergoes adaptations associated with enhanced cocaine locomotor sensitization and place preference (21). In the current study, we tested the hypothesis that cocaine abstinence enhances D2-MSN enkephalin, which auto-inhibits GABA release from these same cells, thereby disinhibiting postsynaptic VP neurons and D1-MSNs to facilitate cocaine seeking.

## Materials and Methods

Male and female mice with conditional deletion of proenkephalin (*Penk*, D2-PenkKO), MORs from D2-MSNs (D2-MORKO), or MORs from D1-MSNs (D1-MORKO), together with littermate and *Adora2a-Cre* controls, were used to examine the role of striatal enkephalin signaling in cocaine-induced circuit plasticity and behavior. Experimental approaches included *ex vivo* electrophysiology, operant self-administration of cocaine and sucrose pellets, quantitative PCR, immunohistochemistry, and RNAscope *in situ* hybridization. Detailed descriptions of animals, surgical procedures, behavioral paradigms, electrophysiological recordings, molecular assays, histology, and statistical analyses are provided in the Supplemental Information.

## Results

### Deletion of enkephalin from D2-MSNs prevents GABA suppression in the striatopallidal circuit during cocaine abstinence

D2-PenkKO mice and *Adora2a*-*Cre* controls received repeated cocaine (15 mg/kg, IP x 10 days) followed by forced abstinence (13-22 days) (Figure 1A-C). Striatal slices were then prepared, and GABA release was measured from ChR2-expressing D2-MSNs onto fluorescently identified VP neurons that project to the VTA (mCherry-positive, Figure 1D). After confirming a synaptic connection between D2-MSNs and the recorded VP neuron via oIPSCs in normal aCSF, a Ca^2+^-free, Sr^2+^-containing aCSF was used to measure aIPSCs. This approach allows us to assess the basal synaptic state by measuring GABA release events selectively arising from D2-MSNs 50–500 ms after D2-MSN optogenetic stimulation (Figure 1E) (13,22–25).

**Figure 1.**
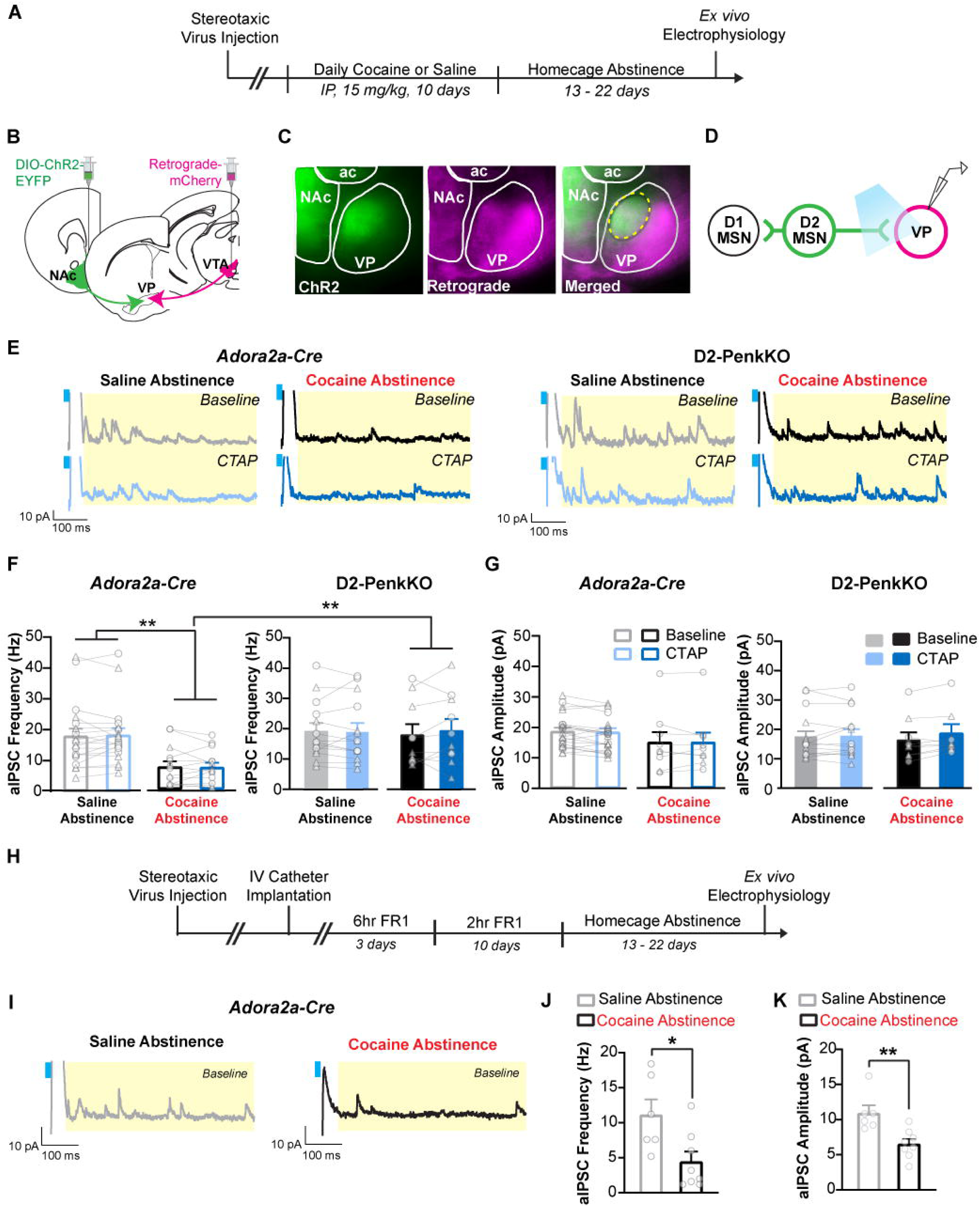
Striatal enkephalin is necessary for cocaine abstinence to suppress GABA release from D2-MSNs to the VP. (A) Experiment timeline of *ex vivo* electrophysiology following forced abstinence from experimenter-administered cocaine injections. (B) Schematic of bilateral AAV infusion. Cre-dependent, EYFP-tagged channelrhodopsin (DIO-ChR2-EYFP) was infused into the nucleus accumbens (NAc), and retrograde AAV-mCherry was infused into the ventral tegmental area (VTA). (C) Representative brain slice images showing ChR2-EYFP and mCherry fluorescence converging in the ventral pallidum (VP, dashed yellow circle). (D) Schematic of *ex vivo* electrophysiology recordings in the VP. D2-MSN terminals were optogenetically stimulated, and recordings were performed in mCherry-positive VP neurons. (E) Representative traces of asynchronous inhibitory postsynaptic currents (aIPSC). The yellow box denotes the 50-550 ms analysis window after optogenetic stimulation (cyan). (F) Cocaine abstinence (black bars) suppressed aIPSC frequency compared with saline abstinence (grey bars) in *Adora2a-Cre* controls (saline: 19 cells/10 mice, cocaine: 10 cells/5 mice), but not in D2-PenkKO mice (saline: 14 cells/5 mice, cocaine: 10 cells/5 mice). Bath application of CTAP (1 µM, blue bars) did not affect aIPSC frequency. (G) aIPSC amplitude was not affected by cocaine abstinence, deletion of striatal enkephalin, or CTAP. (H) Experiment timeline of *ex vivo* electrophysiology recordings performed in the VP of *Adora2a*-*Cre* mice following abstinence from cocaine self-administration or yoked-saline exposure. (I) Representative traces of aIPSC. (J) Abstinence from cocaine self-administration (black) blunted aIPSC frequency compared to yoked saline-abstinent subjects (grey) (cocaine: 8 cells/3 mice, yoked-saline: 6 cells/2 mice). Mean daily cocaine intake: 29.3 mg/kg. (K) aIPSC amplitude was suppressed by cocaine abstinence. Data are shown as mean ± SEM, with individual data points labeled as male (circles) and females (triangles). * *p* < 0.05, ** *p* < 0.01

GABA release was inhibited, as indicated by reduced aIPSC frequency, depending on cocaine-treatment history and genotype (Genotype x Cocaine: F_1, 49_ = 5.19, *p* = 0.027; Figure 1F). Cocaine abstinence inhibited GABA release in *Adora2a*-*Cre* controls compared with saline-abstinent counterparts (t_49_ = 3.42, *p* = 0.001) and cocaine-abstinent D2-PenkKOs (t_49_ = 3.19, *p* = 0.003) (Figure 1F). Conversely, cocaine abstinence had no effect on D2-PenkKOs, and GABA release was similar between saline-abstinent D2PenkKOs and *Adora2a*-*Cre* controls. The mean amplitude of GABA release events was not affected by cocaine-abstinence in either genotype, suggesting unaltered post-synaptic GABA_A_ receptor function (Figure 1G). Importantly, this GABA plasticity was not limited to experimenter-delivered cocaine. Following abstinence from cocaine self-administration, there was a similar reduction in GABA release in *Adora2a*-*Cre* compared with yoked saline controls (t_12_ = 2.66, *p* = 0.021; Figure 1H-J), and GABA amplitude was modestly inhibited following abstinence from cocaine self-administration (t_12_ = 3.5, *p* = 0.004; Figure 1K). These data indicate striatopallidal GABA plasticity emerges after both passive and voluntary cocaine intake and is present during drug abstinence when the risk of cocaine seeking is heightened.

To evaluate the enduring nature of enkephalin-mediated GABA plasticity, we applied the MOR antagonist CTAP to slices. CTAP did not reverse suppression of GABA release in cocaine-abstinent *Adora2a*-*Cre* controls or alter GABA release in any other group (Figure 1F, G). Similarly, the DOR antagonist naltrindole failed to reverse GABA inhibition (Cocaine: F_1, 12_ = 22.84, *p* = 0.0004) and did not affect aIPSC amplitude (Figure S2). Since *Gi*-coupled GABA_B_ receptors in D2-MSNs autoinhibit GABA release (26), we tested whether GABA_B_ receptors maintained GABA plasticity. However, a GABA_B_ antagonist (CGP 55845), alone or combined with CTAP, did not reverse GABA suppression (Genotype x Cocaine: F_1, 14_ = 7.08, *p* = 0.019; Figure S3).

Since mice lacking D2 receptors in D2-MSNs show increased striatal GABA release (27), we examined whether D2-PenkKOs have decreased D2 receptor expression and function, which could account for the lack of effect of cocaine abstinence on GABA release in these mice. *Drd2* mRNA levels were similar between genotypes and cocaine treatment groups (Figure S4B). Additionally, D2 receptor function, inferred by the ability of the D2-like agonist quinpirole to inhibit D2-MSN GABA release, was similar between genotypes (Quinpirole x Cocaine: F_3,121_ = 2.91, *p* = 0.037; Saline-abstinent, baseline vs. 0.25 µM: t_127_ = 10.44, *p* < 0.0001, baseline vs. 0.5 µM: t_127_ = 13.03, *p* < 0.0001, baseline vs. 1 µM: t_127_ = 13.8, *p* < 0.0001; Figure S4D-G). Following cocaine abstinence, quinpirole was less effective, regardless of genotype (saline vs. cocaine, 0.25 µM: t_171_ = 2.33, *p* < 0.05, 0.5 µM: t_171_ = 2.32, *p* < 0.05, 1 µM: t_171_ = 2.86, *p* < 0.01).

Thus, while cocaine appears to desensitize D2 receptors, as previously reported (28,29), D2 receptor expression and function are not altered in D2-PenkKO mice and therefore cannot explain the absence of GABA plasticity in cocaine-abstinent D2-PenkKO mice.

### Cocaine abstinence does not affect intrastriatal GABA release

In addition to innervating the VP, D2-MSNs make extensive intrastriatal connections with D1-MSNs (30–32). Therefore, we examined whether cocaine abstinence similarly inhibits GABA release from D2-MSNs within this intrastriatal circuit. Recordings of aIPSCs were made from ChR2-negative cells (putative D1-MSNs) in the NAc following forced abstinence from repeated cocaine exposure (Figure 2A-D). In contrast to the VP, cocaine abstinence did not inhibit GABA release onto putative D1-MSNs in either genotype (Figure 2F). Additionally, GABA release was not affected by CTAP application, indicating that cocaine did not increase functional opioid peptide tone in the intrastriatal circuit. The amplitude of GABA responses was also not affected by cocaine abstinence or genotype (Figure 2G).

**Figure 2.**
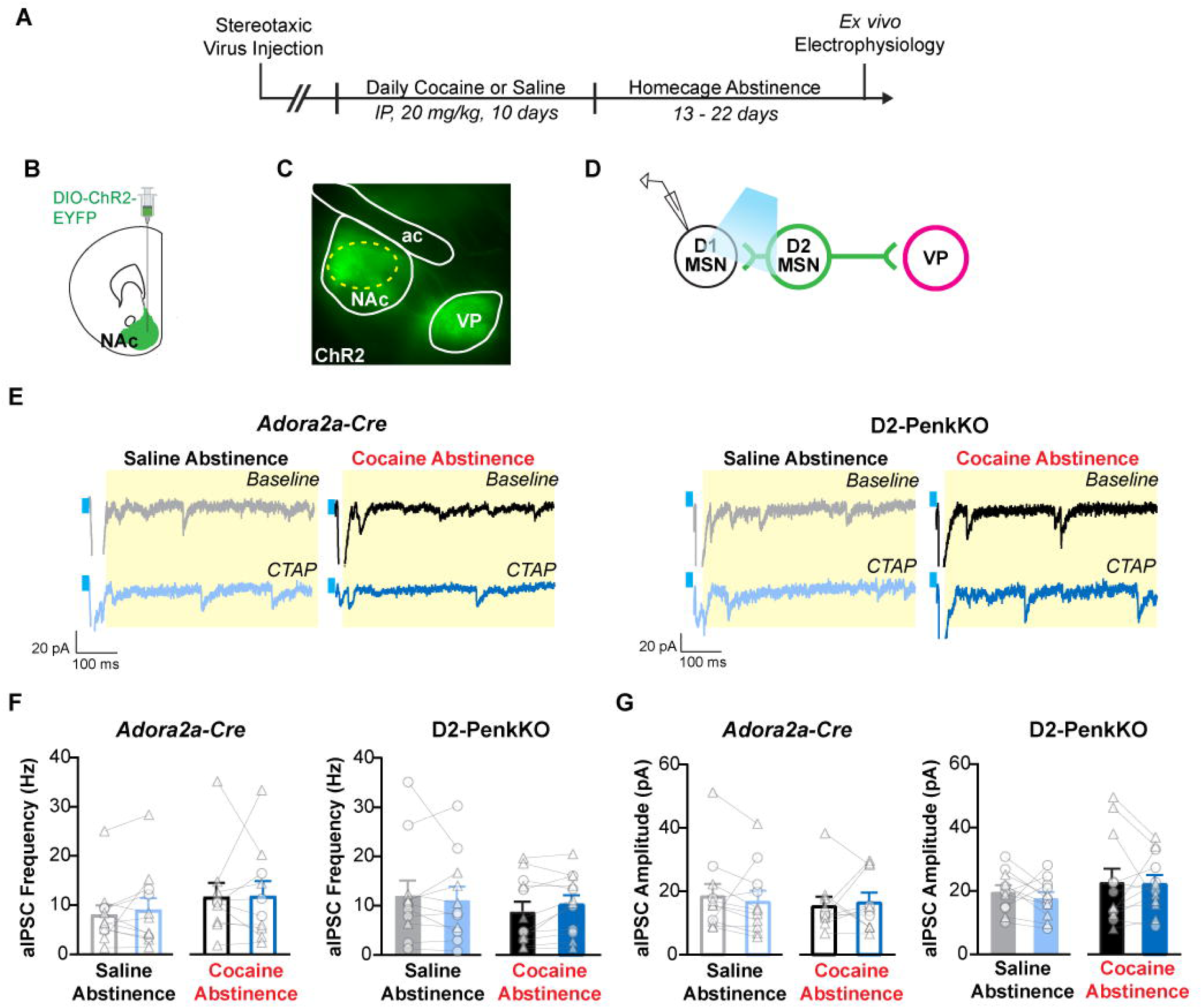
Cocaine abstinence does not inhibit collateral GABA release from D2-MSNs to D1-MSNs. (A) Experiment timeline. (B) Schematic of bilateral AAV-DIO-ChR2-EYFP infusion into the nucleus accumbens (NAc). (C) Representative brain slice image showing ChR2-EYFP fluorescence in the NAc and region of recording (dashed yellow circle). (D) Schematic of *ex vivo* electrophysiology recordings in D1-MSNs. D2-MSN collaterals were optogenetically stimulated, and recordings were performed in putative D1-MSNs (EYFP-negative). (E) Representative traces of asynchronous inhibitory postsynaptic currents (aIPSC). The yellow box denotes the 50-550 ms analysis window after optogenetic stimulation (cyan). (F) Cocaine abstinence (black bars) did not affect aIPSC frequency compared to saline abstinence (grey bars) in *Adora2a-Cre* controls (saline: 11 cells/4 mice, cocaine: 10 cells/4 mice) and D2-PenkKO mice (saline: 11 cells/4 mice, cocaine: 12 cells/4 mice). Bath application of CTAP (1 µM, blue bars) did not affect aIPSC frequency. (G) aIPSC amplitude was not affected by cocaine abstinence, deletion of striatal enkephalin, or CTAP. Data are shown as mean ± SEM, with individual data points labeled as male (circles) and females (triangles). ** *p* < 0.01.

### Cocaine abstinence occludes enkephalin-mediated inhibition of striatopallidal GABA

Our findings thus far suggest that striatal enkephalin is required for the induction, but not maintenance, of striatopallidal GABA plasticity because the plasticity failed to develop in D2-PenkKOs and was not reversed by MOR antagonism in *Adora2a-Cre* controls. We therefore tested whether acute met-enkephalin application suppresses GABA release onto VP neurons. Since exogenous met-enkephalin inhibits GABA release from D2-MSNs onto D1-MSNs (17), we predicted it would have a similar effect in the VP. Consistent with our previous findings, cocaine abstinence inhibited GABA release in the VP of *Adora2a*-*Cre* controls, but not D2-PenkKOs (Genotype x Cocaine x Opioid Treatment: F_2, 70_ = 3.95, *p* = 0.024; *Adora2a*-*Cre*, saline vs cocaine: t_15.43_ = 3.13, *p* = 0.0067; Figure 3D). Met-enkephalin inhibited GABA release in saline-abstinent *Adora2a*-*Cre* controls; however, this effect was occluded in cocaine-abstinent *Adora2a*-*Cre* (saline *Adora2a*-*Cre,* baseline vs Met-ENK: t_10_ = 5.90, *p* = 0.0005; cocaine *Adora2a*-*Cre,* baseline vs Met-ENK: not significant). CTAP reversed the met-enkephalin inhibition in saline-abstinent *Adora2a-Cre* controls (Met-ENK vs CTAP: t_10_ = 5.60, *p* = 0.0007) and marginally increased GABA release in cocaine-abstinent *Adora2a*-*Cre* controls (Met-ENK vs CTAP: t_7_ = 4.72, *p* = 0.0064) to a level substantially less than in saline-abstinent *Adora2a-Cre* mice (CTAP *Adora2a*-*Cre*, saline vs cocaine: t_16.94_ = 2.29, *p* = 0.035). In D2-PenkKOs, met-enkephalin inhibited GABA release in both saline-abstinent and cocaine-abstinent subjects (baseline vs Met-ENK, saline-abstinent: t_10_ = 4.98, *p* = 0.014; cocaine-abstinent: t_8_ = 4.77, *p* = 0.024), and this was fully reversed by CTAP (Met-ENK vs CTAP, saline-abstinent: t_10_ = 6.76, *p* = 0.002; cocaine-abstinent: t_8_ = 5.08, *p* = 0.017).

**Figure 3.**
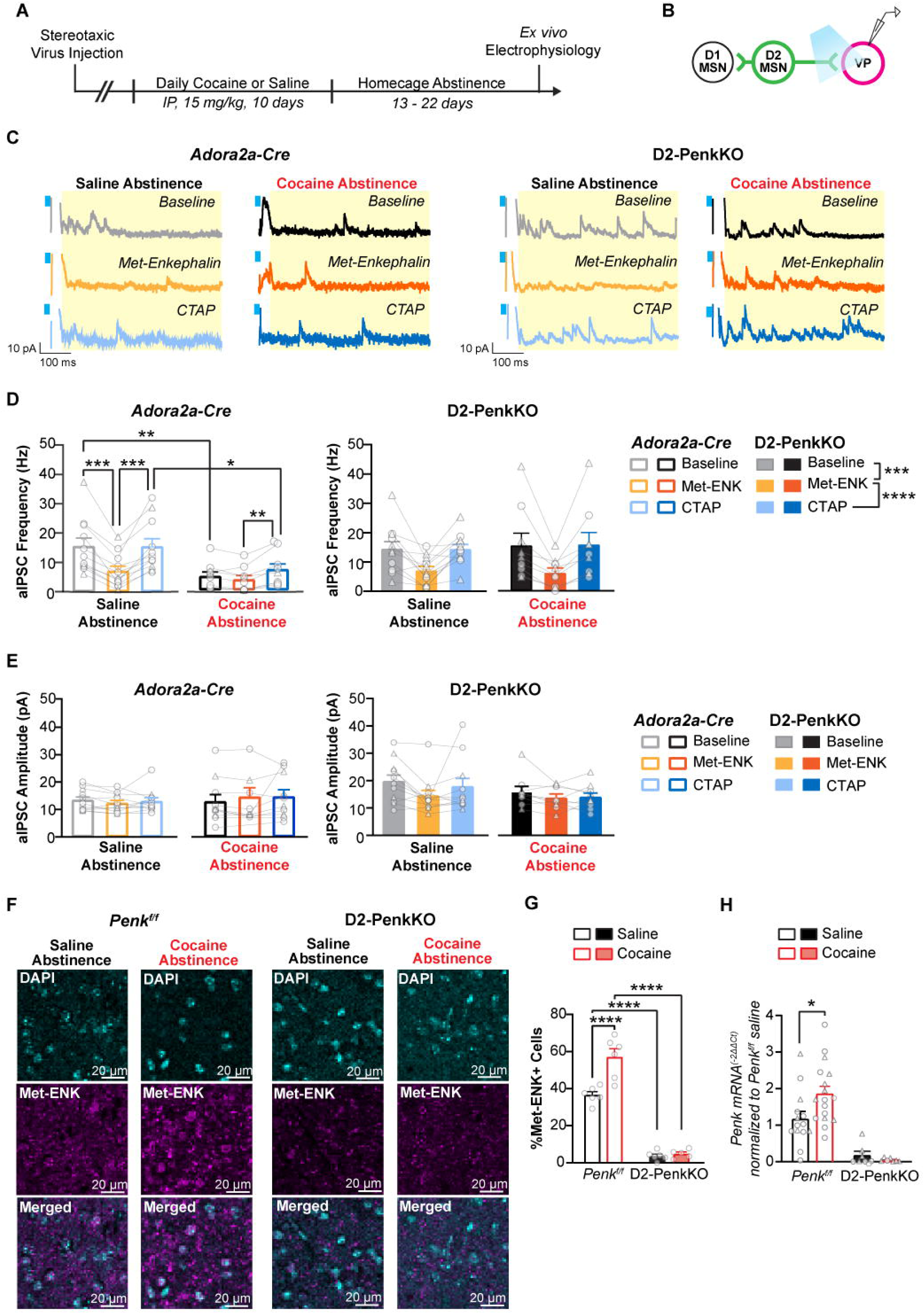
Cocaine-enhanced striatal enkephalin occludes met-enkephalin-induced GABA suppression. (A) Experiment timeline. (B) Schematic of *ex vivo* electrophysiology recordings in the VP. (C) Representative traces of asynchronous inhibitory postsynaptic currents (aIPSC) from saline and cocaine-abstinent *Adora2a-Cre* controls and D2-PenkKOs at baseline and following bath application of met-enkephalin (1 µM, orange) and CTAP (1 µM, blue). The yellow box denotes the 50-550 ms analysis window after optogenetic stimulation (cyan). (D) Cocaine-abstinence inhibited aIPSC frequency compared with saline-abstinence in *Adora2a-Cre* controls (left, saline: 11 cells/4 mice, cocaine: 8 cells/4 mice), but not D2-PenkKOs (saline: 11 cells/4 mice, cocaine: 9 cells/4 mice). Met-enkephalin suppressed aIPSC frequency in saline-abstinent *Adora2a-Cre* and D2-PenkKOs, as well as cocaine-abstinent D2-PenkKOs. CTAP fully reversed this suppression in these groups. CTAP also modestly increased aIPSC frequency in cocaine-abstinent *Adora2a-Cre* controls, but not back to saline levels. (E) aIPSC amplitude was not affected by cocaine abstinence, genotype, or opioid receptor drug application. (F) Representative 40X confocal images of the NAc showing immunohistochemistry fluorescence for met-enkephalin (Met-ENK) positive cells (magenta) in saline-abstinent and cocaine-abstinent D2-PenkKOs and *Penk*^f/f^ controls. (G) Cocaine abstinence (red bars) increased the percentage of met-enkephalin positive cells in the NAc compared with saline abstinence (black bars) for *Penk^f/f^* mice (saline, n = 6; cocaine n = 6) but not D2-PenkKOs (saline, n = 6; cocaine, n = 5). (H) Cocaine abstinence (red bars) increases relative proenkephalin (*Penk*) mRNA abundance in the NAc compared with saline-abstinence (black bars) for control *Penk^f/f^* mice (saline, n = 14; cocaine, n = 17) but not D2-PenkKOs (saline, n = 7; cocaine, n = 7). Data are shown as mean ± SEM, with individual data points labeled as male (circles) and females (triangles). * *p* < 0.05, ** *p* < 0.01, *** *p* < 0.001, **** *p* < 0.0001.

We hypothesized that the lack of effect of met-enkephalin to further inhibit GABA release in cocaine-abstinent *Adora2a-Cre* resulted from occlusion by elevated endogenous enkephalin. In support of this, cocaine abstinence increased striatal *Penk* mRNA (t_29_ = 2.41, *p* = 0.011) and met-enkephalin peptide levels (Genotype x Cocaine: F_1, 19_ = 13.4, *p* = 0.0017; saline vs. cocaine, t_19_ = 5.63, *p* < 0.0001) in *Penk^f/f^* mice (Fig. 3F-H). As shown previously (33), saline-abstinent D2-PenkKOs had extremely low striatal *Penk* mRNA (t_19_ = 3.35, *p* = 0.0034; Fig. 3H) and fewer met-enkephalin positive cells (t_19_ = 9.01, *p* < 0.0001, Fig. 3G) compared with saline-abstinent *Penk^f/f^*controls. Cocaine did not affect these measures in D2-PenkKOs. Collectively, these results indicate that exogenous met-enkephalin suppresses striatopallidal GABA release, but cocaine abstinence occludes this effect due to heightened endogenous enkephalin release from D2-MSNs in control mice.

### Cocaine abstinence gates evoked VP activity through striatal enkephalin

Next, we determined how GABA release from D2-MSNs affects VP neuron excitability and tested whether suppressed GABA release following cocaine abstinence disinhibits VP neuron excitability. A depolarizing current step was applied to VP neurons to elicit approximately 10 action potentials (Baseline, Light Off; Figure 4C, D). The magnitude of the current step was similar between saline-abstinent and cocaine-abstinent mice within each genotype, although D2-PenkKOs required more current than *Adora2a*-*Cre* controls (Genotype: F_1, 29_ = 5.50, *p* = 0.026; Figure S5I). The depolarizing current step was then applied while optogenetically stimulating D2-MSNs (Baseline, Light On; Figure 4D). This procedure was repeated in the presence of met-enkephalin and CTAP.

**Figure 4.**
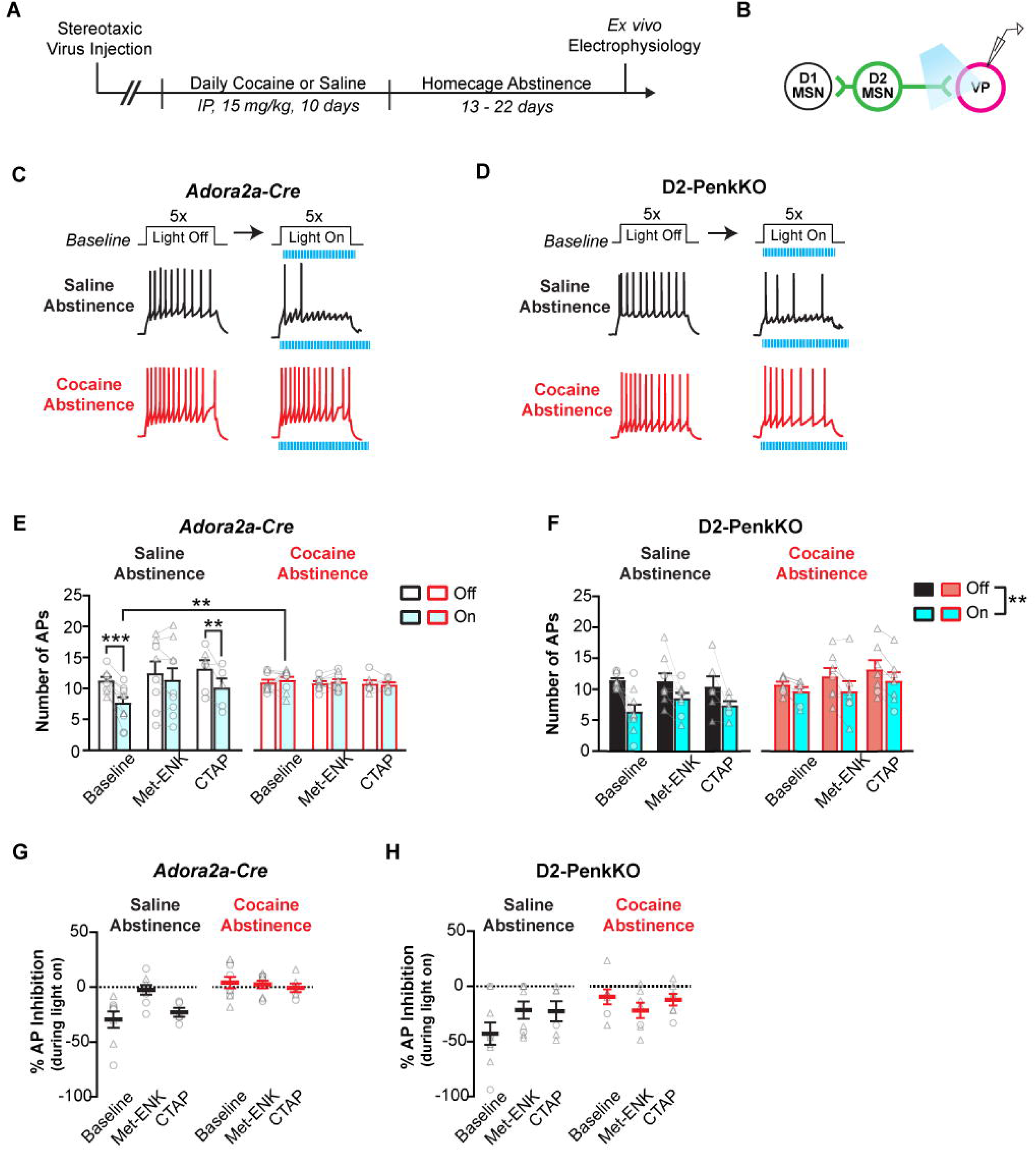
Cocaine abstinence disinhibits VP neurons via the enkephalin-dependent restraint of GABA release from D2-MSNs. (A) Experiment timeline. (B) Schematic of electrophysiology recording in VP neurons. (C, D) Representative evoked action potential (AP) traces for *Adora2a-Cre* controls (C) and D2-PenkKOs (D). A current step was applied to evoke approximately 10 APs and repeated 5 times in absence of D2-MSN optogenetic stimulation (Light Off). The same current step was then applied while simultaneously optogenetically stimulating ChR2-expressing D2-MSN terminals while recording VP firing (Light On). This “Light Off, Light On” protocol was repeated following bath application of met-enkephalin (1 µM) and CTAP (1 µM). (E) In saline-abstinent *Adora2a-Cre* controls (left, n = 8 cells/5 mice), optogenetically-induced GABA release (cyan-filled bars) reduced the number of VP neuron Aps at baseline. This effect was blocked by met-enkephalin and restored by CTAP. Optogenetic stimulation of D2-MSNs did not affect AP firing in cocaine-abstinent *Adora2a-Cre* controls (right, n = 9 cells/5 mice). Asterisks represent post-hoc t-tests (** *p* < 0.01, *** *p* < 0.001). (F) Optogenetic stimulation of D2-MSN terminals suppressed the number of VP APs in saline-abstinent and cocaine-abstinent D2-PenkKOs (saline, n = 9 cells/5 mice; cocaine, n = 7 cells/3 mice). Asterisks represent a main effect of optogenetic stimulation (** *p* < 0.01). (G-H) The effect of optogenetically-evoked GABA release to reduce AP firing is shown as the percent of AP inhibition for *Adora2a-Cre* controls (G) and D2-PenkKOs (H). Data are shown as mean ± SEM, with individual data points labeled as male (circles) and females (triangles).

As predicted, stimulating D2-MSNs reduced VP neuron firing in saline-abstinent *Adora2a*-*Cre* controls (Cocaine x Light x Opioid Treatment: F_2, 19_ = 5.45, *p* = 0.013; Light Off vs Light On: t_7_ = 4.18, *p* = 0.004; Figure 4E). This reduction was blocked by met-enkephalin and restored by CTAP (Light Off vs Light On: t_4_ = 4.56, *p* = 0.01). Conversely, in cocaine-abstinent *Adora2a*-*Cre* controls, D2-MSN stimulation failed to inhibit VP neuron firing, and neither met-enkephalin nor CTAP affected VP neuron excitability (Figure 4E,G). These results indicate that, in saline-abstinent controls, GABA release from D2-MSNs restrains VP neuron excitability, and exogenous met-enkephalin relieves this inhibition. Following cocaine abstinence, however, met-enkephalin no longer disinhibited VP neuron excitability, consistent with occlusion by increased endogenous enkephalin release. Accordingly, D2-MSN stimulation inhibited VP neuron firing in D2-PenkKOs, regardless of cocaine history or opioid drug application (Light: F_1, 14_ = 16.15, *p* = 0.001; Figure 4F,H).

Even though cocaine abstinence disinhibited evoked VP neuron firing by relieving inhibitory input, spontaneous VP firing was suppressed (Genotype x Cocaine: F_1, 31_ = 7.50, *p* = 0.01; saline vs cocaine: t_31_ = 4.11, *p* = 0.0003; Figure S5B, C). This effect appeared to stem from a depolarized action potential threshold (Genotype x Cocaine: F_1, 29_ = 5.30, *p* = 0.029; saline vs. cocaine: t_29_ = 2.90, *p* = 0.0071; Figure S5E and Supplemental Results).

### Striatal enkephalin supports cocaine taking and seeking in females via MORs in D2-MSNs

Because enkephalin is required for abstinence-induced striatopallidal GABA plasticity (Figures 1, 3), we examined whether striatal enkephalin contributes to cocaine taking and cue-induced reinstatement of cocaine seeking. Early in acquisition (FR1), both males and females acquired cocaine self-administration (Lever Presses/hr, Sex x Lever x Trial: F_2.83,113.11_ = 3.60, *p* = 0.018; Figure 5B, E); however, females self-administered more than males (Intake, Sex x Trial: F_3.09,123.7_ = 3.90, *p* = 0.009; Figure 5C, F), and D2-PenkKOs lever-pressed less than controls (Genotype: F_1,40_ = 6.60, *p* = 0.014). Follow-up analysis within each sex separately indicated D2-PenkKO females exhibited reduced lever pressing (Genotype: F_1,21_ = 4.57, *p* = 0.04; Figure 5C) and cocaine intake (Genotype: F_1,18_ = 6.05, *p* = 0.024; Figure 5F) relative to controls. Later in acquisition (FR3), intake in D2-PenkKO females normalized to control levels, suggesting striatal enkephalin supports early acquisition, but not maintenance, of cocaine taking in females. Consistent with this, after establishing stable self-administration neither genotype nor sex affected cocaine taking when tested in dose response, in which all subjects exhibited comparable inverted-U shaped dose-response curves across a range of cocaine doses (Dose: F_1.72,61.04_ = 34.29, *p* < 0.0001; Figure S6I-L).

**Figure 5.**
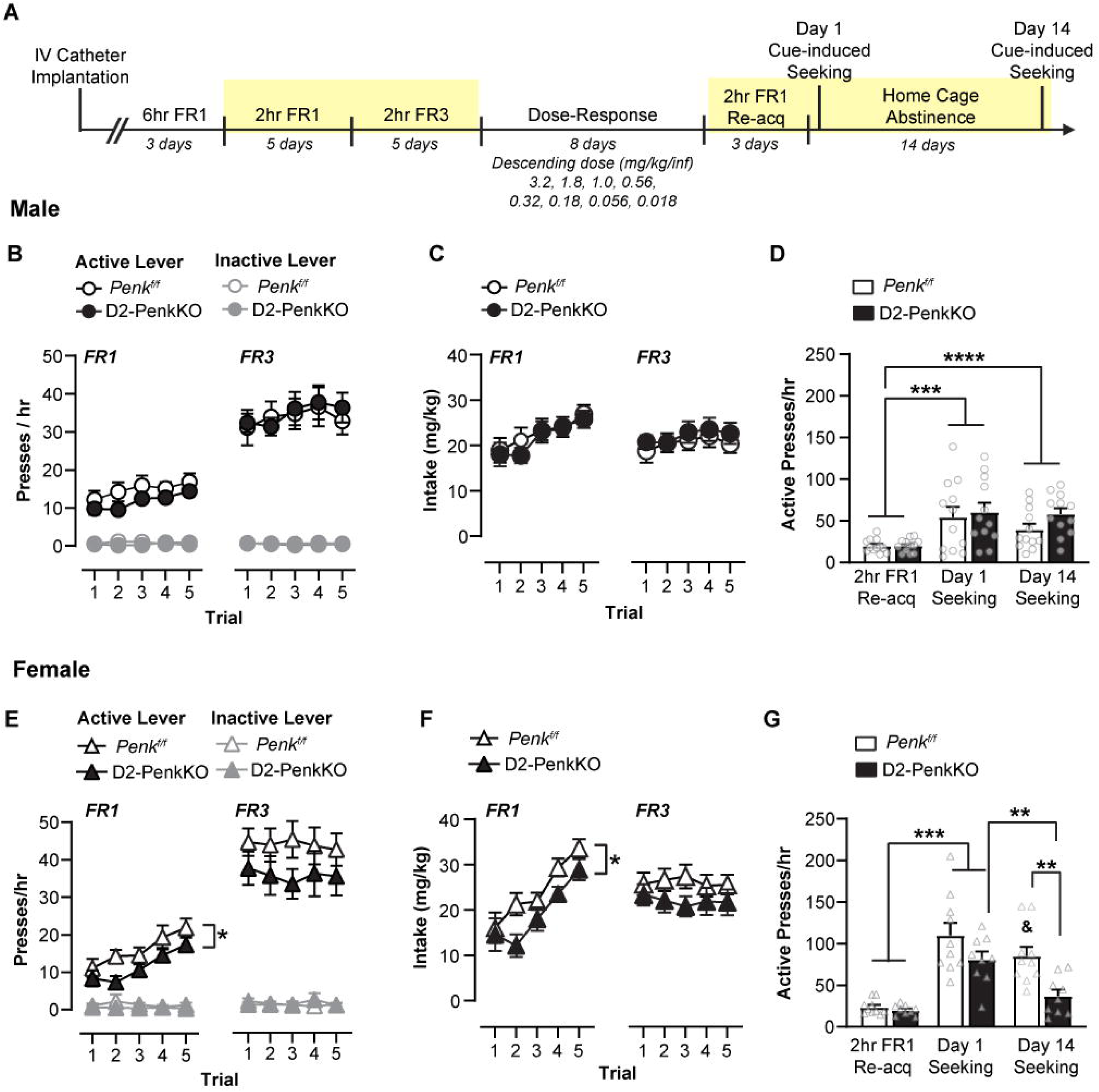
Striatal enkephalin supports early acquisition of cocaine self-administration and cue-indued reinstatement of cocaine seeking in a sex-dependent manner. (A) Experimental timeline for cocaine self-administration in D2-PenkKOs and littermate controls. The yellow box denotes the experimental phases to which the displayed data correspond. (B, E) Active (dark symbols) and inactive lever (light symbols) presses per hour over 10 daily training trials in male (B) and female (E) D2-PenkKOs (filled, male n = 13; female n = 9) and *Penk^f/f^*littermate controls (open, male n = 12; female n = 11). (C, F) Daily total cocaine intake in male (C) and female (F). (D, G) Average active lever presses per hour are shown for the last 2 days of re-acquisition and cue-induced seeking tests performed after 1 and 14 days of forced abstinence from cocaine self-administration in males (D) and females (G). Data are shown as mean ± SEM, with individual data points labeled as male (circles) and females (triangles). *: *p* < 0.05, **: *p* < 0.01, *** *p* < 0.001, **** *p* < 0.0001, & *p <* 0.001 Re-acq vs. Day 14.

We performed additional experiments in D2-MORKOs to examine whether MORs expressed in D2-MSNs mediate some of enkephalin’s effects on cocaine self-administration (Figure 6A). Although both male and female D2-MORKOs acquired cocaine self-administration, (Trial x Lever: F_4.70,159.64_ = 11.83, *p* < 0.001; Figure 6D), female D2-MORKOs exhibited reduced active lever pressing over the last three days of acquisition compared to controls (Sex x Genotype: F_1, 34_ = 6.56, *p* = 0.015; female D2-MORKO vs MOR^f/f^: t_34_ = 2.54, *p* = 0.016; Figure 6E). This genotype effect was not observed in males (Figure 6B-C). Conversely, both male and female D1-MORKOs showed lever discrimination and response rates similar to controls during FR1 and FR3 training (FR1: Trial x Lever: F_4,92_ = 3.28, *p* < 0.015; FR3: Trial x Lever: F_2.38,75.98_ = 3.28, *p* < 0.022; Figure 6G-J), though females exhibited slightly greater lever pressing than males at FR1 (Sex: F_1,23_ = 5.40, *p* = 0.029). These data indicate that MOR signaling in D2-MSNs contributes to the early acquisition phase of cocaine self-administration in females, whereas MORs in D1-MSNs are not required for this early operant learning.

**Figure 6.**
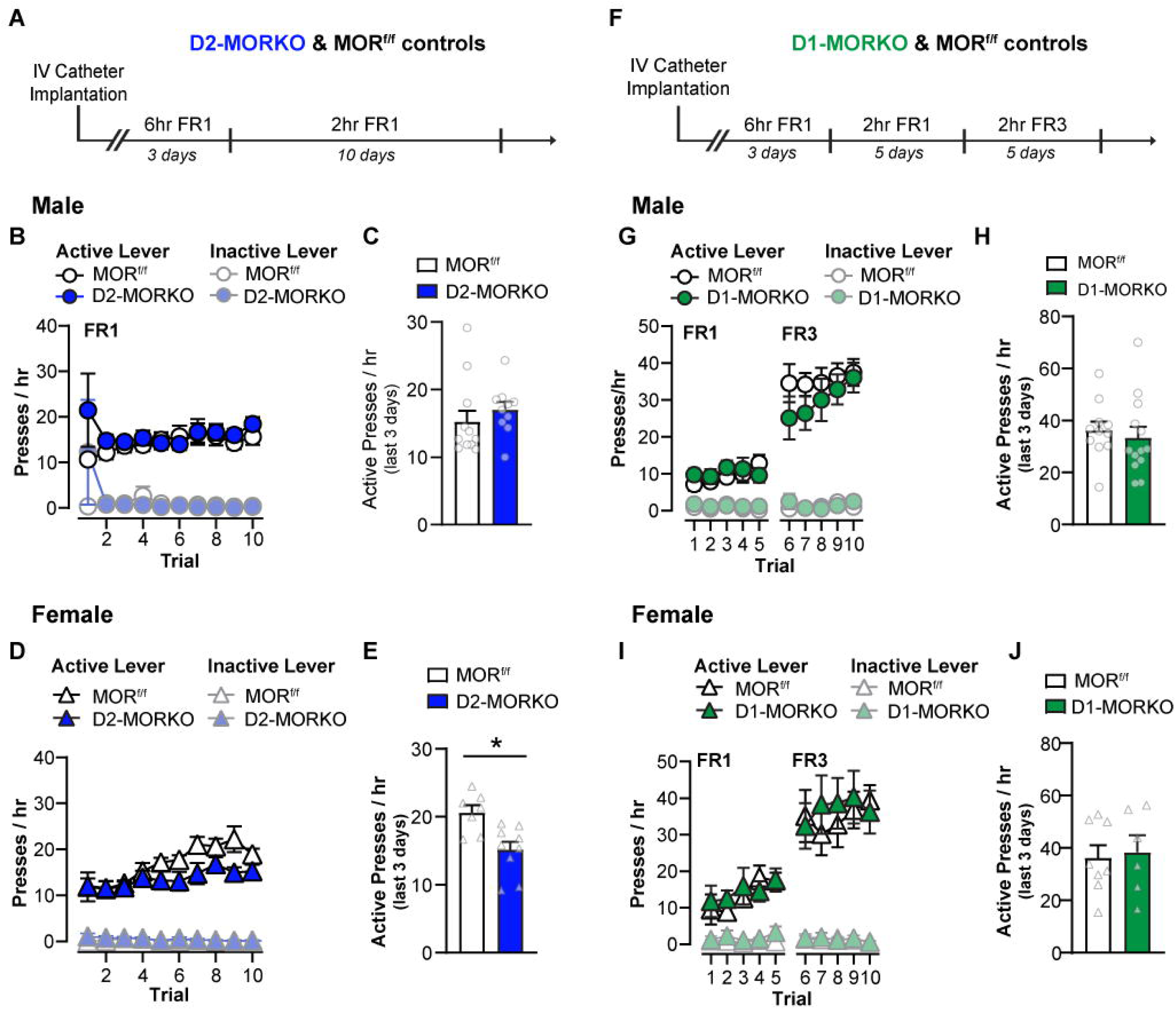
Mu-opioid receptors expressed in D2-MSNs facilitate operant responding for cocaine selectively in females. (A) Experimental timeline for cocaine self-administration in D2-MORKOs and littermate controls. (B, D) Active (dark symbols) and inactive lever (light symbols) presses per hour over 10 daily training trials in male (B) and female (D) D2-MORKOs (blue filled, male n = 10; female n = 10) and MOR^f/f^ littermate controls (black open, male n = 12; female n = 7). (C, E) Average active lever presses over the last 3 days of training were decreased in female D2-MORKOs compared to controls, but not in male D2-MORKOs. (F) Experimental timeline for cocaine self-administration in D1-MORKOs and littermate controls. (G, I) Active (dark symbols) and inactive lever (light symbols) presses per hour over 10 daily training trials in male (G) and female (I) D1-MORKOs (green filled, male, n = 11; female, n = 6) and MOR^f/f^ littermate controls (black open, male, n = 11; female, n = 8). (H, J) Average active lever presses over the last 3 days did not differ between genotypes for males (H) or females (J). ** *p* < 0.01 Data are shown as mean ± SEM, with individual data points labeled as male (circles) and females (triangles).

In examining cue-induced cocaine seeking in D2-PenkKOs following forced abstinence, we found that female D2-PenkKOs exhibited reduced seeking (Genotype x Sex x Trial: F_1.66,63.13_ = 5.19, *p* = 0.012; Figure 5G). Conversely, males, regardless of genotype, maintained cocaine seeking after 1 and 14 days of abstinence compared with re-acquisition of baseline responding at FR1 (Re-acq) when cocaine was available (Re-acq vs Day 1: t_22_ = 4.90, *p* = 0.0002; Re-acq vs Day 14: t_22_ = 5.99, *p* < 0.0001; Figure 5D). Similarly, female *Penk*^f/f^ controls showed robust cocaine seeking across both tests compared with baseline (Re-acq vs Day 1: t_9_ = 6.24, *p* = 0.0005; Re-acq vs Day 14: t_9_ = 5.54, *p* = 0.001) and greater seeking relative to males on day 1 (Day 1, male vs female: t_18.5_ = 2.87, *p* = 0.01). Conversely, female D2-PenkKO mice only exhibited cocaine seeking on day 1 (t_8_ = 6.62, *p* = 0.0005). After 14 days of abstinence, seeking was substantially reduced (D2-PenkKO vs *Penk*^f/f^: t_15.8_ = 3.57, *p* = 0.0026) and no longer greater than baseline (D2-PenkKO: Re-acq vs Day 14, not significant; Figure 5G).

### Striatal enkephalin deletion does not affect sucrose pellet self-administration

We examined whether striatal enkephalin facilitates operant self-administration sucrose pellets (Figure 7A). Mice showed similar lever pressing in early acquisition (FR1-FR3; Trial x Lever: F_1.76,59.91_ = 25.71, *p* < 0.001; Figure 7B, D) and earned more pellets across training regardless of sex and genotype (Trial: F_1.96, 66.71_ = 6.13, *p* = 0.004; Figure 7C, E). Additionally, although female D2-PenkKOs earned nominally fewer pellets over the first three days of FR1 compared with controls (t_18_ = 2.00, *p* = 0.06; Figure 7H), consumption was comparable between genotype and sex (Figure 7G, I). There were also no genotype or sex differences in lever presses or rewards per hour at higher FRs (FR5 – FR32), and D2-PenkKOs did not exhibit altered economic demand for sucrose pellets (see Supplemental Results and Figure S6A-G). In tests of cue-induced sucrose seeking following abstinence, all mice, regardless of sex or genotype, exhibited increased active lever pressing on the first day of seeking relative to acquisition at FR1 (Day: F_2, 61_ = 140.2, *p* < 0.0001; FR1 Acq vs Day 1 Seeking: t_61_ = 16.48, *p* < 0.0001; Figure 7J, K). Seeking remained elevated after 14 days of sucrose abstinence (t_61_ = 5.04, *p* < 0.0001) but had begun to decline (Day 1 vs 14: t_61_ = 10.26, *p* < 0.0001). Thus, striatal enkephalin is not required for the self-administration, consumption, or cue-induced seeking of a sucrose reward.

**Figure 7.**
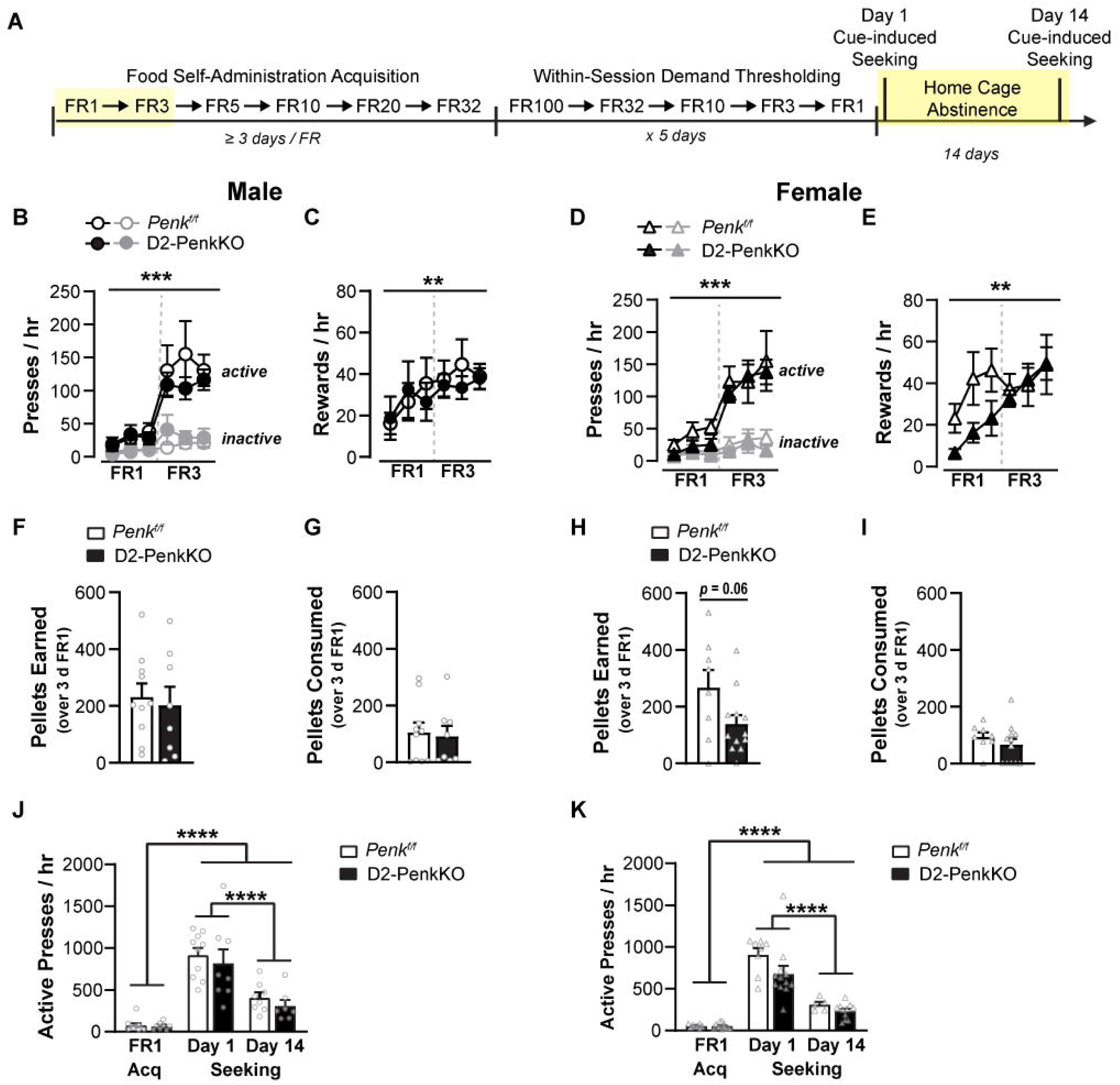
Striatal enkephalin deletion does not affect sucrose pellet self-administration. (A) Experimental timeline for sucrose pellet self-administration in D2-PenkKOs and littermate controls. The yellow box denotes the experimental phases to which the displayed data correspond. Remaining data are shown in Figure S6. (B, F) Active (black) and inactive lever (grey) presses per hour are shown for male (B) and female (F) D2-PenkKOs (filled symbols, male, n = 6; female, n = 9) and *Penk*^f/f^ littermate controls (open symbols, male, n = 10; female, n = 5). Operant responding for cocaine was decreased during the first three days of FR1 in female D2-PenkKOs compared to same-sex controls. (C, D, G, H) Female (G, H) D2-PenkKOs, but not male (C, D), earned fewer rewards per hour than controls during the first three days of FR1 training. (E, I) Male (E) and female (I) D2-PenkKOs consumed a similar number of sucrose reward pellets than same-sex controls. ** *p* < 0.01, *** *p* < 0.001. Data are shown as mean ± SEM, with individual data points labeled as male (circles) and females (triangles).

## Discussion

Animal models and clinical evidence suggest that the risk of relapse increases with progressive drug abstinence, and that this is accompanied by various neural circuit adaptations. Here, we identified striatal enkephalin as a necessary mechanism gating D2-MSN circuit plasticity, as well as the early acquisition of cocaine self-administration and cue-induced reinstatement of cocaine-seeking in females. Moreover, we show that cocaine induces this plasticity selectively at D2-MSN-to-VP synapses, while sparing intrastriatal synapses onto D1-MSNs. Consequently, D2-MSN-mediated restraint of VP neuron firing is blunted following cocaine abstinence. Evidence from parallel self-administration experiments indicates enkephalin released from D2-MSNs suppresses GABA release from the same cells and contributes to cocaine-seeking behavior selectively in females under reinstatement conditions (Figure 8). These results reveal a novel mechanism whereby cocaine abstinence enhances enkephalin release from D2-MSNs, which shifts striatopallidal circuit function in a manner that enhances cocaine-related behaviors.

**Figure 8.**
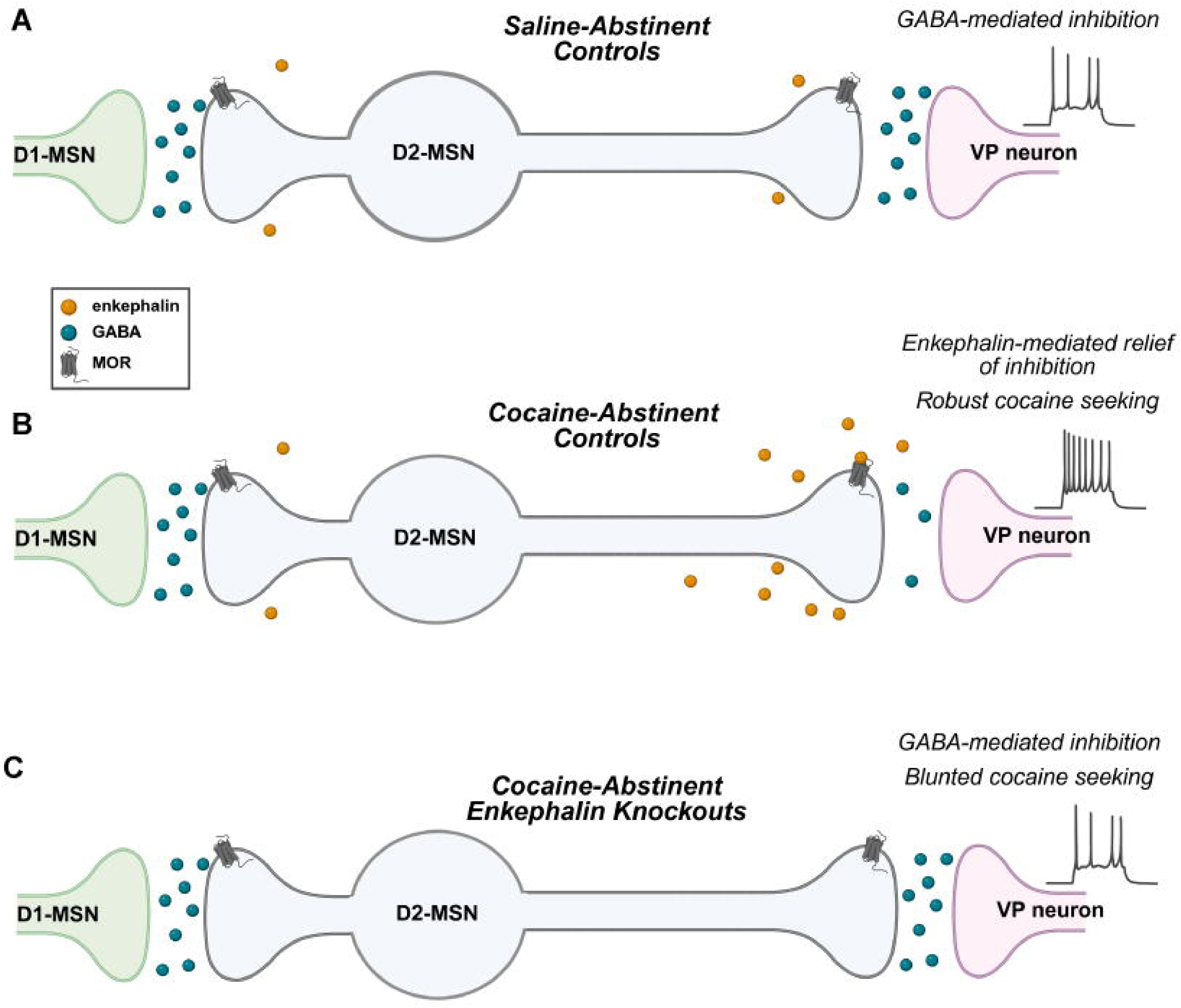
Schematic summarizing the effect of cocaine abstinence on striatal enkephalin, D2-MSN GABA release, VP excitability, and cocaine seeking. (A) In saline-abstinent *Adora2a*-*Cre* controls, GABA released from D2-MSNs inhibits ventral pallidum (VP) neuron firing. (B) In *Adora2a*-*Cre* controls, abstinence from chronic cocaine increases enkephalin from D2-MSNs, which inhibits GABA release from the same cells onto VP neurons, but not onto D1-MSNs, potentially via activation of µ-opioid receptors (MOR) in D2-MSNs. Relief of inhibitory tone onto VP neurons consequently increases their firing. Prolonged cocaine abstinence (14 d) is also associated with increased cue-induced reinstatement of cocaine seeking in females. (C) In mice lacking striatal enkephalin, D2-MSN GABA release is not suppressed following cocaine abstinence. Thus, inhibitory restraint over VP neuron firing is intact. Lack of striatal enkephalin also blunts cue-induced cocaine seeking in females following an abstinence period corresponding to when GABA plasticity and increased seeking are present in controls. Figure Created in BioRender.

### Cocaine-enhanced enkephalin induces persistent GABA suppression in the striatopallidal circuit

Basal GABA release from D2-MSNs to the VP is suppressed following cocaine abstinence, as indicated by fewer asynchronous release events. This is consistent with previous literature that indirectly examined GABA plasticity. High-frequency-stimulation induced long-term depression of GABA in the VP of saline-abstinent, but not cocaine-abstinent subjects (10,12,13). We now show that striatal enkephalin is responsible for this effect. Although enkephalin release was not directly measured, collective evidence from this and other studies (17–19) supports the idea that cocaine abstinence increases enkephalin release from D2-MSNs, which restrains GABA release from the same cells. Consistent with this, bath-applied met-enkephalin was ineffective in further suppressing GABA release in cocaine-abstinent control mice, indicating the effect was occluded.

Suppression of GABA release during cocaine abstinence may represent persistent long-term depression involving structural remodeling of D2-MSN terminals in the VP and recruitment of other receptor systems. Accordingly, GABA suppression during cocaine abstinence was not reversed by opioid or GABA_B_ receptor antagonists. By contrast, GABA suppression induced by bath-applied met-enkephalin was readily reversed by a MOR antagonist, suggesting chronically heightened enkephalin and prolonged MOR/DOR activation may be necessary for lasting GABA plasticity in this circuit.

Other receptors on D2-MSNs, such as those for substance P, nociceptin, and neurokinin1, may contribute to this lasting plasticity. Receptor antagonists for these peptides, however, have previously failed to reverse D2-MSN-to-D1-MSN GABA suppression induced by endogenous opioid peptides (17). The CART peptide (cocaine- and amphetamine-regulated transcript) is another candidate. Its gene (*Cartpt*) is highly expressed in the striatum and upregulated in patients with a history of cocaine use, as well as in mice with elevated striatal *Penk* (34,35). Although the CART receptor has not yet been identified, precluding pharmacological testing, global CART knockout mice may be useful for exploring this further.

Structural remodeling of D2-MSN projections and opioid receptors may also contribute to the enduring nature of GABA suppression following cocaine abstinence. The density of D1-MSN projections into the VP is decreased when striatal D2 receptors are downregulated (21,36,37). It is unknown if similar structural changes occur following cocaine treatment or in D2-MSN terminals, although cocaine decreases striatal D2 receptor availability and function (21,28,29) and D2 receptor activation increases MSN axon collaterals (32). Thus, examining how D2-MSN terminals are remodeled following cocaine abstinence may shed light on the persistence of GABAergic plasticity involving D2-MSNs.

Structural changes may also occur at the receptor level. Reductions in opioid receptor reserve and axonal surface density occur *in vitro* following repeated injections of morphine *in vivo* or prolonged exposure to a selective MOR agonist *in vitro* (38,39). Depleted opioid receptors on D2-MSN terminals due to chronically enhanced enkephalin release would be consistent with our finding that opioid antagonists do not reverse GABA suppression in cocaine-abstinent animals. Alternatively, cocaine-enhanced enkephalin release may have induced constitutive MOR activity in D2-MSNs, similar to withdrawal from repeated morphine administration (40). Consistent with this interpretation, bath application of an inverse MOR agonist increased striatopallidal GABA release in cocaine-abstinent rats, revealing constitutive MOR signaling induced by cocaine exposure (12).

### Striatal enkephalin gates cocaine-induced VP activity and cocaine taking and seeking

Our current-clamp experiments revealed that D2-MSN GABA release restrains evoked VP neuron firing in saline-abstinent, but not cocaine-abstinent, *Adora2a-Cre* mice. This mechanism could account for heightened VP neuron activity during cue-induced seeking trials following extinction from cocaine self-administration (11) and would be consistent with reports that activation of the VP is both necessary and sufficient for cue-induced cocaine seeking (7,8,10). Importantly, this cocaine-induced disinhibition of VP neurons was absent in D2-PenkKOs, in which GABA release remained intact. These findings identify striatal enkephalin as a mediator of cocaine-induced striatopallidal circuit adaptations that may contribute to the disinhibited VP state associated with cocaine seeking. Cocaine abstinence, however, reduced the spontaneous activity of VP neurons. This may in part be explained by heightened GABA release from D1-MSNs to the VP, which has been reported following cocaine abstinence (13). Lower baseline spontaneous activity was associated with a depolarized AP threshold, suggesting cocaine abstinence alters voltage-gated sodium channels in VP neurons. D2-PenkKOs also exhibited lower spontaneous VP neuron firing, suggesting that low striatal enkephalin may recapitulate the effects of cocaine abstinence on circuit function. Loss of striatal enkephalin also appeared to affect susceptibility to further circuit disruption by cocaine, as cocaine abstinence perturbed VP neuron resting membrane potential in D2-PenkKOs but not in *Adora2a*-*Cre* controls.

Although our findings identify striatal enkephalin as a mediator of both striatopallidal plasticity and cocaine-related behaviors, they do not establish that the behavioral effects are directly mediated by this plasticity. Nevertheless, female D2-PenkKOs and D2-MORKOs showed reduced cocaine self-administration during early acquisition, and female D2-PenkKOs exhibited reduced cue-induced cocaine seeking following an abstinence period corresponding to the time when striatopallidal GABA plasticity is present. Moreover, a similar striatopallidal plasticity emerged following abstinence from self-administered cocaine, consistent with the idea that these circuit adaptations contribute to cocaine-seeking behavior during abstinence. Collectively, our findings suggest enkephalin may act through MORs in D2-MSNs to support early acquisition of cocaine self-administration, and that striatopallidal GABA plasticity is associated with cocaine seeking. Whether these same MORs also contribute to cocaine seeking, though, remains unclear because cue-induced reinstatement was not examined in D2-MORKOs.

Finally, we observed sex differences in how striatal enkephalin affects cocaine seeking and taking. Female D2-PenkKOs failed to develop striatopallidal GABA suppression and showed reduced cocaine seeking, suggesting this form of plasticity may be functionally linked to cocaine-seeking behavior in females. Conversely, although D2-PenkKO males also failed to develop striatopallidal GABA suppression, they showed robust cocaine seeking, suggesting that this plasticity is not a major contributor to cocaine seeking in males or that competing adaptations occur. Multiple circuit adaptations likely work in concert to drive cocaine seeking, and our findings suggest the relative contribution of each adaptation differs by sex, with enkephalin-mediated striatopallidal GABA plasticity likely playing a stronger role in shaping circuit adaptations linked to cocaine-seeking behavior in females.

### Conclusions

We have identified striatal enkephalin as a gating mechanism by which abstinence from chronic cocaine alters striatopallidal circuit function to enhance VP excitability. This neuroadaptation is persistent and associated with cue-induced reinstatement of cocaine seeking in females. Our findings also raise the intriguing possibility that synthetic opioids may engage this circuit in a manner analogous to enkephalin, potentially enhancing the vulnerability to engage in opioid-cocaine co-use.

## Supporting information

Supplemental Methods, Figures, and Results

## Author Contributions

KM conceptualized and contributed to electrophysiology and self-administration experiment data collection and data analysis, generated figures and graphics, and wrote the first draft of the paper. PL contributed to electrophysiology data collection. AN contributed to data collection and analysis for knockout validation, self-administration, and qPCR experiments. IBC contributed to knockout validation data collection and analysis. BR, MS, MV, DS, AB, and GG contributed to self-administration data collection and analysis. LKD led study design and conceptualization, supervised the research, provided financial support, and contributed to data analysis, figure generation, and manuscript preparation. All authors reviewed and edited the manuscript.

## Data Availability

Earlier versions of these data were presented at SfN, GRC, and INRC (41–43). This manuscript is also presented as a preprint (44).

## Resources and Other

We would like to acknowledge Dr. Robert Messing, Dr. Laura Colgin, Dr. Hitoshi Morikawa, Dr. Regina Mangieri, Dr. Lief Fenno, and Dr. Caitlin Orsini for their guidance and feedback. We are grateful to Dr. Andreas Zimmer for providing the floxed *Penk* mice. We also would like to acknowledge the dedicated UT Austin veterinary staff for their support in maintaining our mouse colony.

## References

1. Parvaz MA, Moeller SJ, Goldstein RZ (2016): Incubation of Cue-Induced Craving in Adults Addicted to Cocaine Measured by Electroencephalography. JAMA Psychiatry 73: 1127– 1134.

2. Li X, Venniro M, Shaham Y (2016): Translational Research on Incubation of Cocaine Craving. JAMA Psychiatry 73: 1115–1116.

3. Zhao D, Zhang M, Tian W, Cao X, Yin L, Liu Y, et al. (2021): Neurophysiological correlate of incubation of craving in individuals with methamphetamine use disorder. Molecular Psychiatry 2021 26:11 26: 6198–6208.

4. Scheyer AF, Loweth JA, Christian DT, Uejima J, Rabei R, Le T, et al. (2016): AMPA Receptor Plasticity in Accumbens Core Contributes to Incubation of Methamphetamine Craving. Biol Psychiatry 80: 661–670.

5. Krasnova IN, Marchant NJ, Ladenheim B, Mccoy MT, Panlilio L V, Bossert JM, et al. (2008): Incubation of Methamphetamine and Palatable Food Craving after Punishment-Induced Abstinence. Neuropsychopharmacology 39. 10.1038/npp.2014.50

6. Venniro M, Zhang M, Shaham Y, Caprioli D (2017): Incubation of Methamphetamine but not Heroin Craving After Voluntary Abstinence in Male and Female Rats. Neuropsychopharmacology 42: 1126–1135.

7. Mahler S V, Vazey EM, Beckley JT, Keistler CR, Mcglinchey EM, Kaufling J, et al. (2014): Designer receptors show role for ventral pallidum input to ventral tegmental area in cocaine seeking. Nat Neurosci 17: 577–585.

8. Farrell MR, Ruiz CM, Castillo E, Faget L, Khanbijian C, Liu S, et al. (2019): Ventral pallidum is essential for cocaine relapse after voluntary abstinence in rats. Neuropsychopharmacology 44: 2174–2185.

9. Smith KS, Tindell AJ, Aldridge JW, Berridge KC (2009): Ventral pallidum roles in reward and motivation. Behavioural Brain Research 196: 155–167.

10. Heinsbroek JA, Neuhofer DN, Griffin WC, Siegel GS, Bobadilla AC, Kupchik YM, Kalivas PW (2017): Loss of plasticity in the D2-accumbens pallidal pathway promotes cocaine seeking. Journal of Neuroscience 37: 757–767.

11. Heinsbroek JA, Bobadilla AC, Dereschewitz E, Assali A, Chalhoub RM, Cowan CW, Kalivas PW (2020): Opposing Regulation of Cocaine Seeking by Glutamate and GABA Neurons in the Ventral Pallidum. Cell Rep 30: 2018–2027.e3.

12. Kupchik YM, Scofield MD, Rice KC, Cheng K, Roques BP, Kalivas PW (2014): Cocaine dysregulates opioid gating of GABA neurotransmission in the ventral pallidum. Journal of Neuroscience 34: 1057–1066.

13. Creed M, Ntamati NR, Chandra R, Lobo MK, Lüscher C (2016): Convergence of Reinforcing and Anhedonic Cocaine Effects in the Ventral Pallidum. Neuron 92: 214–226.

14. Gerfen CR, Surmeier DJ (2011): Modulation of striatal projection systems by dopamine. Annu Rev Neurosci 34: 441–466.

15. Fricker LD, Margolis EB, Gomes I, Devi LA (2020): Five decades of research on opioid peptides: Current knowledge and unanswered questions. Mol Pharmacol 98: 96–108.

16. Banghart MR, Neufeld SQ, Wong NC, Sabatini BL (2015): Enkephalin Disinhibits Mu Opioid Receptor-Rich Striatal Patches via Delta Opioid Receptors. Neuron 88: 1227–1239.

17. Dai KZ, Choi IB, Levitt R, Blegen MB, Kaplan AR, Matsui A, et al. (2022): Dopamine D2 receptors bidirectionally regulate striatal enkephalin expression: Implications for cocaine reward. Cell Rep 40: 111440.

18. Crespo JA, Manzanares J, Oliva JM, Corchero J, Palomo T, Ambrosio E (2001): Extinction of cocaine self-administration produces a differential time-related regulation of proenkephalin gene expression in rat brain. Neuropsychopharmacology 25: 185–194.

19. Mongi-Bragato B, Zamponi E, García-Keller C, Assis MA, Virgolini MB, Mascó DH, et al. (2016): Enkephalin is essential for the molecular and behavioral expression of cocaine sensitization. Addiction Biology 21: 326–338.

20. Mongi-Bragato B, Avalos MP, Guzmán AS, Bollati FA, Cancela LM, Toll L, et al. (2018): Enkephalin as a pivotal player in neuroadaptations related to psychostimulant addiction. Front Psychiatry 9: 1–11.

21. Dobbs LK, Kaplan AR, Bock R, Phamluong K, Shin JH, Bocarsly ME, et al. (2019): D1 receptor hypersensitivity in mice with low striatal D2 receptors facilitates select cocaine behaviors. Neuropsychopharmacology 44: 805–816.

22. Kircher DM, Aziz HC, Mangieri RA, Morrisett RA (2019): Ethanol experience enhances glutamatergic ventral hippocampal inputs to D1 receptor-expressing medium spiny neurons in the nucleus accumbens shell. Journal of Neuroscience 39: 2459–2469.

23. Bekkers JM, Clements JD (1999): Quantal amplitude and quantal variance of strontium-induced asynchronous EPSCs in rat dentate granule neurons. Journal of Physiology 516: 227–248.

24. Xu-Friedman MA, Regehr WG (2000): Probing fundamental aspects of synaptic transmission with strontium. Journal of Neuroscience 20: 4414–4422.

25. Gerdeman G, Lovinger DM (2001): CB1 cannabinoid receptor inhibits synaptic release of glutamate in rat dorsolateral striatum. J Neurophysiol 85: 468–471.

26. Dobbs LK, Kaplan AR, Lemos JC, Matsui A, Rubinstein M, Alvarez VA (2016): Dopamine Regulation of Lateral Inhibition between Striatal Neurons Gates the Stimulant Actions of Cocaine. Neuron 90: 1100–1113.

27. Lemos JC, Friend DM, Kaplan AR, Shin JH, Rubinstein M, Kravitz A V., Alvarez VA (2016): Enhanced GABA Transmission Drives Bradykinesia Following Loss of Dopamine D2 Receptor Signaling. Neuron 90: 824–838.

28. Gong S, Fayette N, Heinsbroek JA, Ford CP (2021): Cocaine shifts dopamine D2 receptor sensitivity to gate conditioned behaviors. Neuron 109: 3421–3435.e5.

29. Nader MA, Morgan D, Gage HD, Nader SH, Calhoun TL, Buchheimer N, et al. (2006): PET imaging of dopamine D2 receptors during chronic cocaine self-administration in monkeys. Nature Neuroscience 2006 9:8 9: 1050–1056.

30. Wilson CJ, Groves PM (1980): Fine structure and synaptic connections of the common spiny neuron of the rat neostriatum: A study employing intracellular injection of horseradish peroxidase. Journal of Comparative Neurology 194: 599–615.

31. Taverna S, Ilijic E, Surmeier DJ (2008): Recurrent Collateral Connections of Striatal Medium Spiny Neurons Are Disrupted in Models of Parkinson’s Disease. Journal of Neuroscience 28: 5504–5512.

32. Lalchandani RR, Vicini S (2013): Inhibitory collaterals in genetically identified medium spiny neurons in mouse primary corticostriatal cultures. Physiol Rep 1: 1–12.

33. Matsumura K, Nicot A, Choi IB, Asokan M, Le NN, Natividad LA, Dobbs LK (2023): Endogenous opioid system modulates conditioned cocaine reward in a sex-dependent manner. Addiction Biology 28. 10.1111/ADB.13328

34. Guerri L, Dobbs LK, da Silva e Silva DA, Meyers A, Ge A, Lecaj L, et al. (2023): Low Dopamine D2 Receptor Expression Drives Gene Networks Related to GABA, cAMP, Growth and Neuroinflammation in Striatal Indirect Pathway Neurons. Biological Psychiatry Global Open Science 3: 1104–1115.

35. Ong ZY, McNally GP (2020): CART in energy balance and drug addiction: Current insights and mechanisms. Brain Res 1740: 146852.

36. Inbar K, Levi LA, Kupchik YM (2022): Cocaine induces input and cell-type-specific synaptic plasticity in ventral pallidum-projecting nucleus accumbens medium spiny neurons. Neuropsychopharmacology 47: 1461–1472.

37. Cazorla M, deCarvalho FD, Chohan MO, Shegda M, Chuhma N, Rayport S, et al. (2014): Dopamine D2 Receptors Regulate the Anatomical and Functional Balance of Basal Ganglia Circuitry. Neuron 81: 153–164.

38. Fyfe LW, Cleary DR, Macey TA, Morgan MM, Ingram SL (2010): Tolerance to the antinociceptive effect of morphine in the absence of short-term presynaptic desensitization in rat periaqueductal gray neurons. Journal of Pharmacology and Experimental Therapeutics 335: 674–680.

39. Jullié D, Benitez C, Knight TA, Simic MS, von Zastrow M (2022): Endocytic trafficking determines cellular tolerance of presynaptic opioid signaling. Elife 11. 10.7554/eLife.81298

40. Wang D, Raehal KM, Lin ET, Lowery JJ, Kieffer BL, Bilsky EJ, Sadée W (2004): Basal Signaling Activity of μ Opioid Receptor in Mouse Brain: Role in Narcotic Dependence. J Pharmacol Exp Ther 308: 512–520.

41. Matsumura K, Dobbs LK (2025): Striatal enkephalin gates cocaine-induced GABA plasticity and excitability of the ventral pallidum. Society for Neuroscience. San Diego.

42. Matsumura K, Dobbs LK (2026): Striatal enkephalin gates disinhibition of ventral pallidum neurons after cocaine abstinence and supports cocaine seeking. Gordon Research Conference: Neurobiology of Addiction. Barcelona.

43. Matsumura K, Dobbs LK (2026): Striatal enkephalin gates disinhibition of ventral pallidum neurons after cocaine abstinence and supports cocaine seeking. International Narcotics Research Conference. Denver.

44. Matsumura K, Lyuboslavsky P, Nicot A, Remmers B, Smith M, Choi IB, et al. (2026): Cocaine-Enhanced Enkephalin Gates Plasticity of Distinct Striatal Synapses and Facilitates Cue-Induced Cocaine-Seeking During Abstinence. bioRxiv 2026.03.11.711212.

