## Supplemental Methods, Figures, and Results for "Cocaine-Enhanced Enkephalin Gates Plasticity of Distinct Striatal Synapses and Facilitates Cue-Induced Cocaine-Seeking During Abstinence"

**Supplemental Information**

**Supplemental Methods**

**Animals.** All animal procedures were performed in accordance with guidelines from the University of Texas at Austin Institutional Animal Care and Use Committee. Male and female adult mice with a homozygous deletion of proenkephalin (*Penk*) from D2-MSNs (D2-PenkKO), mu-opioid receptors (MORs) from D2-MSNs (D2-MORKO), or MORs from D1-MSNs (D1-MORKO) were used. Validation of the D2-PenkKO (1) and D2-MORKO (2) lines is described in prior publications, and validation of the D1-MORKO line can be found in the supplemental results (Figure S1). Mice lacking proenkephalin (*Penk*) from D2-MSNs (D2-PenkKO) (1) and MORs from D2-MSNs (D2-MORKO) (2) were generated by crossing *Penk^f/f^* mice (3) or MOR^f/f^ mice (4) with *Adora2a-Cre^+/-^*mice (B6.FVB(Cg)-Tg(Adora2a-cre)KG139Gsat/Mmucd; RRID: MMRRC_036158-UCD). Mice lacking MORs from D1-MSNs were generated by crossing MOR^f/f^ mice with *Drd1-Cre^+/-^* mice (B6.FVB(Cg)-Tg(Drd1-cre)EY217Gsat/Mmucd; RRID: MMRRC_034258-UCD). *Adora2a-Cre^+/-^* mice were used as controls in electrophysiology experiments, and Cre-negative littermates (*Penk^f/f^* and MOR^f/f^) were used as controls in qPCR, immunohistochemistry (IHC), and self-administration experiments. Mice were group-housed in a temperature- and humidity-controlled environment under a 12:12h light cycle with food and water available *ad libitum*, except as described in the self-administration experiments.

**Stereotaxic surgery.** Male and female *Adora2a-Cre^+/-^* and D2-PenkKO mice (8-12 weeks old) were head-fixed on a stereotaxic frame under isoflurane anesthesia and received bilateral AAV infusions (100 nL/min) via a 30-gauge needle attached to a Hamilton Syringe. For VP recordings, mice received Cre-dependent channelrhodopsin (ChR2-EYFP, 300 nL/side) in the NAc shell (AP +1.6, ML ± 0.6, DV -4.5mm) and retrograde AAV-mCherry (200 nL/side) in the ventral tegmental area (VTA; AP -3.15, ML ± 0.4, DV -4.75mm) to identify and record from VP neurons that receive D2-MSN GABA projections and themselves project to the VTA. We focused on the NAc shell 🡪 VP 🡪 VTA circuit because cocaine abstinence inhibits D2-MSN GABA release in this circuit (5) and it is implicated in cocaine seeking (6). Recordings of intrastriatal GABA release from D2-MSNs onto neighboring D1-MSNs were accomplished by infusing ChR2-EYFP into the NAc core (AP +1.3, ML ±1.0, DV -4.6mm). We previously found that GABA release in this NAc core circuit is associated with behavioral responses to cocaine (7). Cre-dependent channelrhodopsin (ChR2) was from Addgene (pAAV-EF1a-double floxed-hChR2(H134R)-EYFP-WPRE-HGHpA, 20298-AAV5, RRID: Addgene_20298, titer: 7.5 x 10^12^). Retrograde AAV-mCherry was from Addgene (rgAAV.hSyn.mCherry, 114472-AAVrg, RRID: Addgene_114472, titer: 9.5 x 10^12^, 200 nL/side). Experiments were performed at least 2 weeks after surgery.

**Catheter surgery.** Male and female adult mice (8 – 23-week-old) were implanted with a catheter in the right jugular vein as previously described (8). A catheter (Instech, C20PU-MJV2012 or C20PU-MJV2013) was inserted into the vein through a small hole and secured with silk suture. The catheter inlet was fed subcutaneously and attached to a vascular access button with 25-gauge pin-port (Instech, VABM1B/25), which was secured between the scapulae. Mice recovered for one week before beginning self-administration.

**Drugs**. Intraperitoneally administered saline (10 mL/kg) or cocaine HCl (Sigma-Aldrich, C5776) dissolved in sterile saline (0.9%) was administered for 10 days (15 or 20 mg/kg) and followed by 14 days of forced abstinence before performing electrophysiology recordings, immunohistochemistry, or quantitative polymerase chain reaction. These doses were chosen because they increase *Penk* mRNA and enkephalin peptide levels in the striatum and increase locomotion (7,9–11). A range of cocaine unit doses (3.2, 1.8, 1.0, 0.56, 0.32, 0.18, 0.056, 0.018) were used in the self-administration experiments (see below for details). Met-enkephalin and naltrindole were from Sigma-Aldrich (M6638, N115), CTAP and quinpirole were from Tocris (1560, 1061), and CGP 55845 HCl was from HelloBio (HB0960).

***Ex vivo* electrophysiology.**

**Slice and experiment preparation**. Following forced abstinence from 10 days of cocaine treatment (IP: 15 mg/kg and 20 mg/kg for VP and D1-MSN recordings, respectively. Self-administration: 1 mg/kg/infusion), mice were transcardially perfused with ice-cold cutting solution and then rapidly decapitated. Sagittal slices (260 µm) were prepared in carbogenated (95%:5% O_2_:CO_2_) cutting solution (in mM: 225 sucrose, 119 NaCl, 26.2 NaHCO_3_, 1 NaH_2_PO_4_, 1.25 D-glucose, 2.5 KCl, 4.9 MgCl_2_, 0.1 CaCl_2_, 3 kynurenic acid) on a vibratome (Campden, 7000-smz). Slices rested for 30 minutes at 32°C and were stored in carbogenated artificial cerebrospinal fluid (aCSF, in mM: 124 NaCl, 1 NaH_2_PO_4_, 2.5 KCl, 1.3 MgCl_2_, 2.5 CaCl_2_, 20 D-glucose, 26.2 NaHCO_3_, 0.4 ascorbate) at 20°C until recording at 31 – 33°C. Slices were visualized using an Olympus BX51 fluorescence microscope equipped with a 40X objective. Whole-cell recordings were made from mCherry-positive VP neurons or putative D1-MSNs (EYFP-negative) using glass electrodes (2 - 4 MΩ) filled with a twice-filtered Cs-based internal solution for voltage-clamp recordings and K^+^-based internal for current-clamp recordings. The internal solution for voltage-clamp recordings in VP neurons contained (in mM): 120 CsMeS, 10 CsCl, 10 HEPES, 0.2 EGTA, 10 p-creatine, 4 Na_2_-ATP, 0.4 NA_2_-GTP). For voltage-clamp recordings in D1-MSNs, the internal solution contained (in mM): 60 CsMeS, 60 CsCl, 10 HEPES, 0.2 EGTA, 10 p-creatine, 4 Na_2_-ATP, 0.4 Na_2_-GTP. For current-clamp recordings, internal solution contained (in mM): 115 KMeSO_4_, 10 KCl, 1.5 MgCl_2_, 10 HEPES, 0.2 EGTA, 10 p-creatine, 4 Na_2_ATP, 0.4 Na_2_GTP. All internal solutions had a pH of 7.2 – 7.3 and an osmolarity of 298 – 300 mosmol/L.

Synaptic responses recorded in D1-MSNs and action potentials recorded in VP neurons were made in the presence of NMDA, AMPA, and GABA_B_ receptor antagonists (5 µM NBQX, 5 µM R-CPP, 2 µM CGP 55845, respectively). Additionally, at the beginning of each voltage-clamp and current-clamp experiment, the synaptic connectivity between D2-MSN terminals and the patched VP neuron or D1-MSN was confirmed by measuring the optogenetically-evoked IPSC (oIPSC). Light was delivered through a fiber optic (473 nm LED, 4 ms; 200 µm/0.22 NA, ThorLabs), and light power was adjusted to evoke a stable oIPSC (200 – 600 pA) over at least 10 minutes before voltage-clamp and current-clamp experiments were performed. For all experiments, cells with input resistance ≥ 1 GΩ, holding current < -200pA, and access resistance ≥ 25 MΩ or that varied > 20% from baseline were excluded. Data were acquired using Multiclamp 700B (Molecular Devices), filtered at 1 kHz, digitized at 10 kHz, and analyzed using pClamp (Clampfit, v.11).

**Voltage-clamp.** To determine whether cocaine abstinence decreases GABA release from D2-MSNs onto VP neurons and D1-MSNs, recordings of asynchronous GABA_A_-mediated inhibitory postsynaptic currents (aIPSC) were made in aCSF in which Ca^2+^ was substituted with strontium chloride (Sigma-Aldrich, 439665; in mM: 124 NaCl, 1 NaH_2_PO4, 2.5 KCl, 1.8 MgCl_2_, 20 d-glucose, 26.2 NaHCO_3_, 0.4 ascorbate, 2 SrCl_2_). The Sr^2+^-aCSF in recordings of VP neurons also contained the nicotinic acetylcholine receptor antagonist DHβE (1 µM, Tocris, 2349) to isolate GABA_A_ synaptic responses. The frequency and amplitude of quantal GABA release events from D2-MSNs were measured 50–550 ms after optogenetic stimulation (5,12–15). VP neurons and D1-MSNs were held at -5 mV and -55 mV, respectively. Recordings were made for 10 min at baseline and during drug washes: met-enkephalin (1 µM), CTAP (1 µM), naltrindole (10 µM), and CGP 55845 HCl + CTAP (2 µM and 1 µM, respectively). Traces were analyzed using template search (Clampfit). To probe D2 receptor sensitivity, recordings of optogenetic IPSCs (oIPSCs) were made from putative D1-MSNs in the NAc core. oIPSCs were measured for 10 minutes at baseline and over increasing concentrations of the D2-like agonist quinpirole (0.25 µM, 0.5 µM, 1 µM,). In both the aIPSC and D2 receptor experiments, after drug washes were completed, the antagonist gabazine (5 µM, Hellobio, HB0901) was applied to confirm the GABA_A_ receptor identity of the synaptic response.

**Current-clamp**. VP neurons were maintained at -80 mV through constant current injection, and recordings were made in the presence of NMDA, AMPA, and GABA_B_ receptor antagonists. A depolarizing current step (800 ms) was applied to evoke ~10 action potentials (Light Off). The same current step was then applied while simultaneously delivering an optogenetic train to stimulate D2-MSN terminals (Light On; 473 nm LED, 4 ms pulse, 16 Hz, 1260 ms duration). Light Off and Light On phases were repeated five times each, and the average number of action potentials was calculated. For recordings of spontaneous action potentials in VP neurons, constant current injection was turned off, and spikes were recorded over 2 minutes. The action potential threshold was determined by extrapolating the voltage at the time when the voltage’s derivative equaled 20 mV/sec from the phase-plane plot generated in Clampfit. After hyperpolarization was calculated as the voltage difference between the baseline membrane potential and the most negative voltage after the depolarizing current step. Evoked and spontaneous action potentials were detected using threshold analysis. The average resting membrane potential was calculated using the statistics mode in Clampfit. Each parameter was measured at baseline and again 10 minutes after met-enkephalin (1 µM) and CTAP (1 µM) wash.

**Self-administration.**

**Apparatus.** Operant self-administration chambers (MedAssociates; 6 cm W × 17 cm L × 13 cm H) were equipped with a house light, ventilation fan, two retractable levers, and a drug availability LED above the active lever. Cocaine was delivered intravenously via tubing connecting a drug infusion pump (6 µl/s) to the subject’s inlet port via a single-channel fluid swivel mounted on a counterbalanced arm (Instech). For the food self-administration experiment, sucrose pellets (20 mg; Bio-Serv, F07595) were delivered from a magazine into a central hopper located between the two levers upon completion of the fixed ratio on the active lever.

**Animals and general procedures.** Adult male and female D2-PenkKOs, D2-MORKOs, D1-MORKOs, and littermate controls were individually housed on a 12:12 reversed light cycle, and experiments were performed during the dark cycle. Mice were mildly food restricted (95 - 90% of free-feeding weight) for up to 7 days to encourage exploration during acquisition. Mice were habituated in the behavior room for 30 minutes prior to the start of each session. The start of all food and cocaine sessions was signaled by the house light turning off, the levers extending, and illumination of a stimulus light above the active lever to indicate cocaine or food availability. Completion of the operant response resulted in extinguishing the stimulus light during cocaine or sucrose pellet delivery. Inactive lever presses were recorded but had no scheduled consequences. The active lever assignment was counterbalanced between subjects.

**Cocaine.** Mice were trained to self-administer cocaine (1.0 mg/kg/infusion) or received yoked saline infusions, on a fixed ratio 1 (FR1) schedule of reinforcement in 6-hour sessions over 2–3 days until they met acquisition criteria (≥ 10 infusions/day, 2:1 active:inactive lever-press ratio for 2 days). Depending on the experimental objective, mice then underwent self-administration procedures that varied in the response schedule and cocaine dose (Table 1). Sessions ended when the duration was reached or when mice self-administered a maximum cocaine dose (40 mg/kg), whichever occurred first. Catheter patency was confirmed by the loss of the righting reflex within 5 seconds of a brevital sodium injection (10 mg/kg, IV). Catheters were maintained by daily heparin flush (30 unit/mL; 0.02 mL), and mice that failed patency testing at any point were excluded from the study.

**Food.** Mice were habituated to sucrose pellets for 3 days in the home cage before beginning self-administration training in daily 2-hour FR1 sessions. The FR was then progressively increased (FR3, FR5, FR10, FR20, FR32), and subjects were required to meet acquisition criteria before progressing to the next FR (≥ 20 pellets/day, 2:1 active:inactive lever press ratio, for 3 days). After completing FR32, mice were tested over 4 days on a within-session thresholding procedure to generate economic demand curves for food. Each day, the FR response was decreased on a half-log scale (FR100, FR32, FR10, FR3, FR1). A descending FR was used to minimize satiation and food caching early in the session. Each FR was available for 5 minutes followed by a 25-minute timeout period during which the levers retracted, cue-light extinguished, house light turned on, and pellets were not delivered. After completing the thresholding procedure, cue-induced reinstatement of sucrose pellet seeking was measured following 1 day and 14 days of sucrose pellet abstinence. Seeking tests were performed under extinction conditions, during which the cue light was presented but responses were not reinforced.

**Quantitative polymerase chain reaction.** Following 14 days of forced abstinence from cocaine (20 mg/kg, IP x 10 days), male and female D2-PenkKOs and littermate controls were deeply anesthetized with isoflurane, decapitated, and the ventral striatum was dissected on ice. The tissue was homogenized, RNA was extracted (Qiagen), and cDNA was synthesized (BioRad). Relative mRNA abundance of proenkephalin (*Penk*, Mm01212875_m1), dopamine D2 receptor (*Drd2*, Mm00438541_m1), µ-opioid receptor (*Oprm1*, Mm01188089_m1), δ-opioid receptor (*Oprd1*, Mm00443063_m1), and β-actin (*Actb,* Mm01205647_g1) were determined using a TaqMan Gene Expression Assay. All probes were sourced from Thermofisher. A CFX384 Real-Time System was used with the following settings: initial hold at 95 °C for 20 sec, 40 cycles at 95 °C for 1 sec, and 60 °C for 20 sec. Samples were run in triplicate, and negative controls were run in parallel. Relative abundance was calculated using ^ΔΔ^Ct.

**Immunohistochemistry.** Following 14 days of forced abstinence from cocaine (20 mg/kg, IP x 10 days), male and female D2-PenkKOs and littermate controls were transcardially perfused with PBS followed by 4% PFA. Brains were harvested and stored in 4% PFA followed by 30% sucrose for 24 hours before being stored in OCT compound at -80°C. Brains were cryosectioned (30 µm, Leica) into PBS containing 0.02% sodium azide. Floating sections were washed (3 x 15 min in PBS, RT), blocked in 10% normal goat serum (1 hour, RT), and then incubated in a rabbit met-enkephalin primary antibody (1:500, Immunostar, Catalog#: 20065, RRID: AB_572250) in 10% normal goat serum for 18 hours at 4°C. Slices were washed again (PBS 3 x 15 min), incubated in anti-rabbit Alexa Fluor-647 (1:1000, Thermofisher, Catalog#: A-21245, RRID: AB_2535813) in 1% normal goat serum for 2 hours at RT. Slices were washed (3 x 15 min, PBS), mounted, and coverslipped (DAPI Fluoromount, SouthernBiotech). Images of the NAc shell were acquired (3 images/hemisphere) on a confocal microscope (Nikon AXR laser scanning) with a PlanFluor 40x, NA-1.3, WD-0.2 mm oil immersion objective. Met-enkephalin-positive cells were counted (CellProfiler, v.4.2.8) and averaged across 18 images taken from 3 slices per mouse.

**RNA *in situ* hybridization.** D1-MORKO and MOR^f/f^ mice were deeply anesthetized with isoflurane and their brains were removed and flash frozen in isopentane on dry ice. Coronal cryosections (12 um) containing the striatum and periaqueductal grey (PAG; control brain region) were mounted onto slides and fixed in 4% paraformaldehyde in PBS (1hr, 4°C). Sections were rinsed in 1X PBS (2 x 1 min, RT), dehydrated in ethanol (50% x 5 min, 70% x 5 min, 100% 2x 5 min, RT) and stored overnight in 100% ethanol (-20°C). We used the RNAscope Multiplex Fluorescent Reagent Kit v2 (Advance Cell Diagnostics) with a custom-designed *Oprm1* probe (Mm-Oprm1-O4-C2; 544731) to detect MORs and a *Penk* probe (Mm-Penk; 318761) to label striatal D2-MSNs. Sections were treated with H_2_O_2_ (10 min, RT), rinsed (Milli-Q H_2_O, 2 x 1 min, RT), incubated with RNAscope Protease IV (30 min, RT), and rinsed again (Milli-Q H_2_O, 2 x 1 min). Sections were then incubated in amplification buffers 1 and 2 (1 x 30 min each, 40°C), and 3 (1 x 15 min, 40°C), with 1X wash buffer rinses between each amplification buffer (2 x 2 min, RT). Probes were developed individually using HRP-C1 and HRP-C2 (15 min each, 40°C), an opal dye (30 min, 40°C), and HRP blocker (15 min, 40°C). Sections were washed with 1X wash buffer (2 x 2 min, RT) between developing each probe. Sections were cover slipped with DAPI Fluoromount (SouthernBiotech) and images of the ventral striatum and periaqueductal gray were acquired (2-4 images/section, 1-6 sections/region) on a fluorescent microscope (Zeiss) with a 40x oil immersion objective. *Oprm1* puncta were quantified (CellProfiler v4.2.8) in DAPI-positive cells classified as *Penk*-positive (putative D2-MSNs) or *Penk*-negative (non-D2-MSNs, considered putative D1-MSNs). Cells containing at least two *Oprm1* puncta were considered MOR-positive, and the percentage of MOR-positive cells within each population was calculated to assess MOR reduction in putative D2- and D1-MSNs in the striatum and PAG.

**Data Analysis.** Electrophysiology data were analyzed using two- and three-way repeated measures analysis of variance (RM-ANOVA), with genotype, sex, and cocaine treatment as between-subjects factors and drug application as a within-subjects factor. Self-administration of cocaine and food outcome measures were analyzed using two-, three-, and four-way RM-ANOVA or mixed effects analysis, with genotype and sex as between-subjects factors and trial and lever (when appropriate) as within-subjects factors. Demand curves for food self-administration data were constructed by fitting the Hursh-Silberberg exponential demand model (16) using the Excel-based Exponential Demand Calculator developed by Steven Hursh (v4-2, 2011). Food rewards earned were modeled as a function of “price”, defined as the fixed ratio (FR) response requirement. Food rewards earned at each FR were averaged across subjects within each sex and genotype, and demand curves were fit to these group means. Because the exponential model requires log-transformed values, zero values at FR100 were substituted with 0.01 before model fitting. The model was used to estimate Q_0_, the predicted food rewards earned at minimal cost (demand setpoint) and α, which represents demand elasticity (the rate at which rewards earned declines as price increases). Normalized Pmax (nPmax), an index of motivation representing the price at which responding is maximal and associated with progressive ratio breakpoints (17), was calculated as described by Bentzley et al. (2013). Because D2-PenkKOs have lower striatal *Penk* mRNA than littermate controls (1), an unpaired t-test was used to analyze cocaine’s effect on *Penk* mRNA expression within each genotype. Relative *Drd2* mRNA expression and met-enkephalin peptide expression were analyzed using a two-way ANOVA with genotype and cocaine treatment as between-subjects factors. Multifactorial repeated measures datasets with missing values were analyzed using a mixed effects model. Violations of sphericity in repeated measures were corrected using the Greenhouse-Geisser method. Significant interactions were followed up with Šidák-corrected multiple comparisons. Results were considered significant at an alpha of 0.05. Data were analyzed and graphed with GraphPad Prism (v. 11.0.2) and SPSS Statistics (v. 31.0, IBM). Significant main effects and interactions are reported in text. Non-significant main effects and interactions for voltage-clamp (Table 2), current-clamp (Table 3), molecular (Table 4), and behavioral experiments (Table 5) are reported in tables. Data are presented as mean ± standard error of the mean, and individual data points are labeled by sex.

**Supplemental Results**

To validate the deletion of MORs from D1-MSNs in our conditional knockout line, D1-MORKO, we examined *Oprm1* mRNA abundance in the NAc using qPCR (Figure S1A). D1-MORKO mice showed decreased abundance of *Oprm1* mRNA compared with MOR^f/f^ controls (t_10_ = 5.19, *p* = 0.0004; Figure S1A). Abundance of transcript for the delta-opioid receptor (*Oprd1*) or *Penk* was not changed. To confirm this *Oprm1* levels were selectively reduced in D1-MSNs and preserved in D2-MSNs, we examined cell-type specificity using and RNA *in situ* hybridization (Figure S1B-D). Co-expression of *Oprm1* with *Penk* was used to identify MOR-expressing putative D2-MSNs, and *Penk*-negative cells were considered putative D1-MSNs. The percentage of *Oprm1*-positive putative D1-MSNs, but not putative D2-MSNs, was reduced in D1-MORKOs relative to controls (t_21_ = 7.60, *p* < 0.0001; Figure S1C). Additionally, there was no difference between genotypes in the percentage of *Oprm1*-positive cells in the periaqueductal grey, a control region.

Deletion of striatal enkephalin reduced spontaneous VP neuron activity (saline-abstinent *Adora2a*-Cre vs. D2-PenkKO: t_31_ = 3.28, *p* = 0.0026; Figure S5B, C). This reduction was also associated with a depolarized action potential threshold at baseline (saline-abstinent, D2-PenkKO vs. *Adora2a*-Cre: t_29_ = 2.08, *p* = 0.046; Figure S5E) and a hyperpolarized resting membrane potential following cocaine abstinence (Genotype x Cocaine: F_1, 31_ = 4.60, *p* = 0.040; D2-PenkKO saline vs. cocaine: t_31_ = 2.19, *p* = 0.037; cocaine-abstinent, D2-PenkKO vs. Adora2a-Cre: t_31_ = 2.06, *p* = 0.048; Figure S5D). Neither action potential afterhyperpolarization nor input resistance were altered by cocaine or enkephalin deletion (Figure S5F-H).

When tested in a thresholding procedure, D2-PenkKOs and littermate controls similarly decreased the number of sucrose pellets earned as the price (i.e., FR) increased, and this effect was similar between sexes (Price: F_1.4,36.4_ = 36.0, *p* < 0.0001; Figure S6D, G). Parameters estimated from the demand curve including behavioral elasticity (alpha), the basal hedonic setpoint for sucrose pellets (Q_0_), and motivation for sucrose pellets (nPmax) were also similar between genotype and sex (Figure S6D, G, inset). Collectively with the acquisition data, these findings indicate that striatal enkephalin modestly contributes to the early acquisition of palatable food self-administration, but not to the economic demand, motivation, or consumption of a sucrose reward.

**Supplemental Figures**

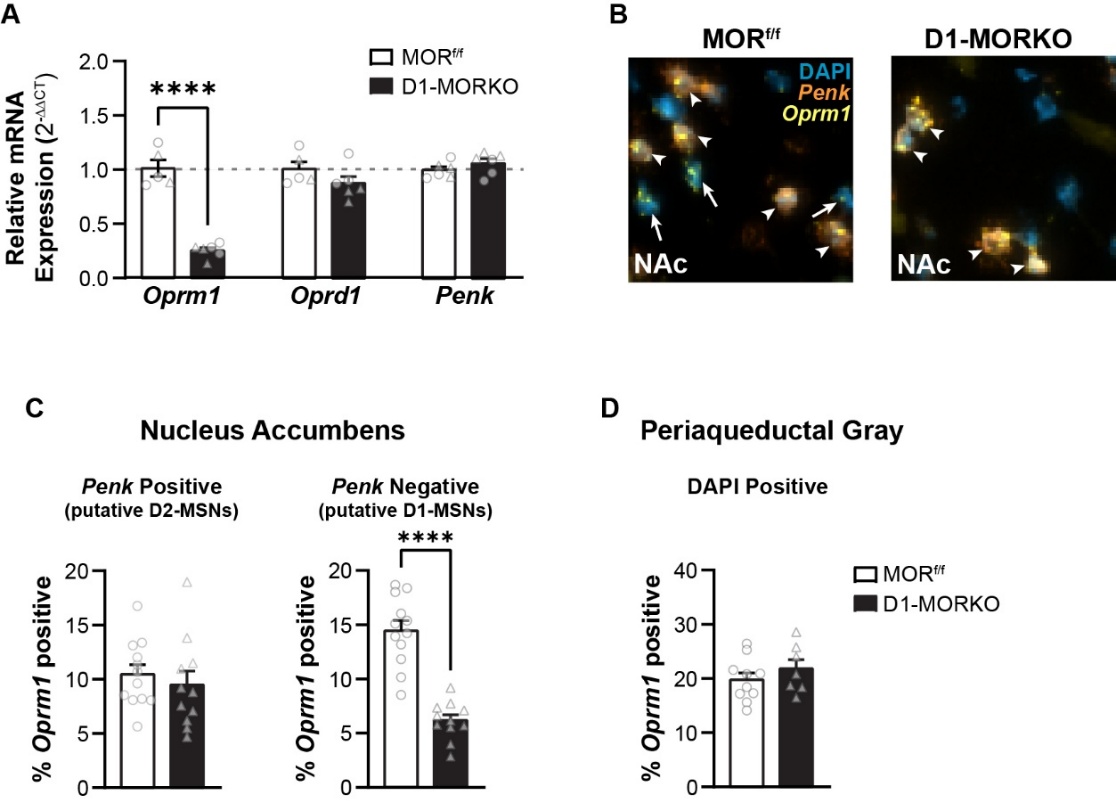

**Supplemental Figure 1.** **D1-MORKO mice have reduced abundance of *Oprm1* mRNA in striatal D1-MSNs.** (A) Relative abundance of *Oprm1* mRNA is decreased in the NAc of D1-MORKO mice (black) compared to littermate controls (white). Abundance of the delta-opioid receptor (*Oprd1*) and proenkephalin (*Penk*) are not altered. (B) Representative RNA *in situ* hybridization images acquired in the NAc are shown for D1-MORKO (right) and MOR^f/f^ littermate controls (left). Co-expression (white arrows) of *Oprm1* (yellow) with *Penk* (orange) indicates the presence of MORs in putative D2-MSNs in both genotypes. *Oprm1* was detected in *Penk*-negative cells (DAPI, cyan, white arrows), which include D1-MSNs, in MOR^f/f^ controls but not D1-MORKOs. (C) The percentage of *Penk*-positive cells (putative D2-MSNs) that express MORs is equal between genotypes (left). In contrast, the percentage of *Penk*-negative cells (putative D1-MSNs) that express MORs is significantly reduced in D1-MORKOs (right). (D) The percentage of MOR-positive cells in the periaqueductal gray, a control region, is similar between genotypes. Data are shown as mean ± SEM, with individual data points labeled as male (circles) and female (triangles). **** *p* < 0.0001.

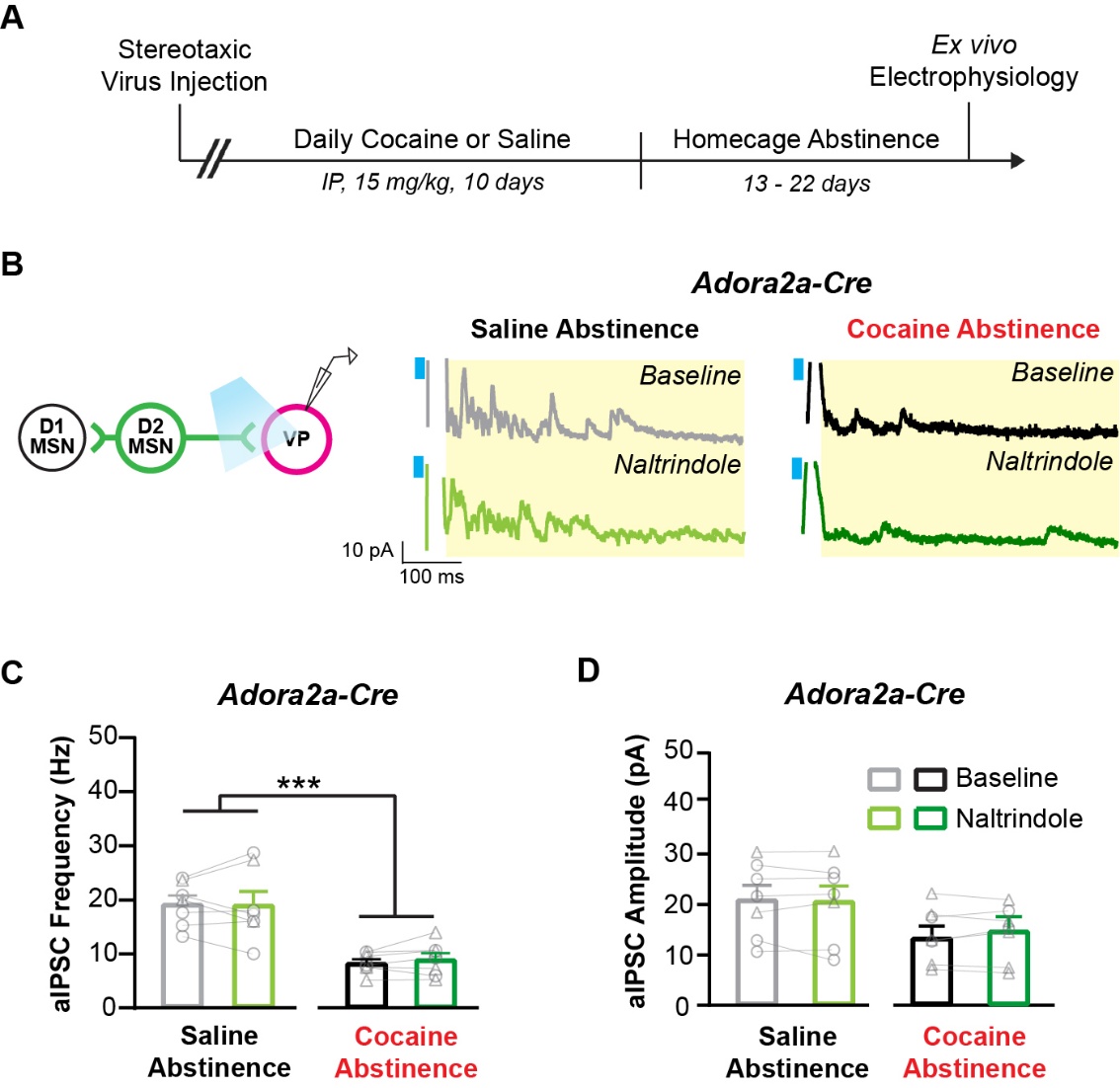

**Supplemental Figure 2.** **A delta opioid receptor antagonist does not reverse cocaine-abstinence-induced, striatal enkephalin-dependent GABA suppression in the ventral pallidum.** (A) Experiment timeline. (B) Electrophysiology recording configuration (left). Representative traces (right) of asynchronous inhibitory postsynaptic currents (aIPSCs) from saline- and cocaine-abstinent *Adora2a*-Cre controls at baseline and following bath application of the DOR antagonist, naltrindole (10 µM). The yellow box denotes the 50 – 500 ms analysis window after optogenetic stimulation (cyan). (C) Cocaine abstinence reduced aIPSC frequency in *Adora2a*-Cre controls compared to saline-treated counterparts (saline: 7 cells/3 mice; cocaine: 7 cells/3 mice). (D) Average aIPSC amplitude was not affected by cocaine abstinence or naltrindole. Data are shown as mean ± SEM, with individual data points labeled as male (circles) and female (triangles). *** *p* < 0.001.

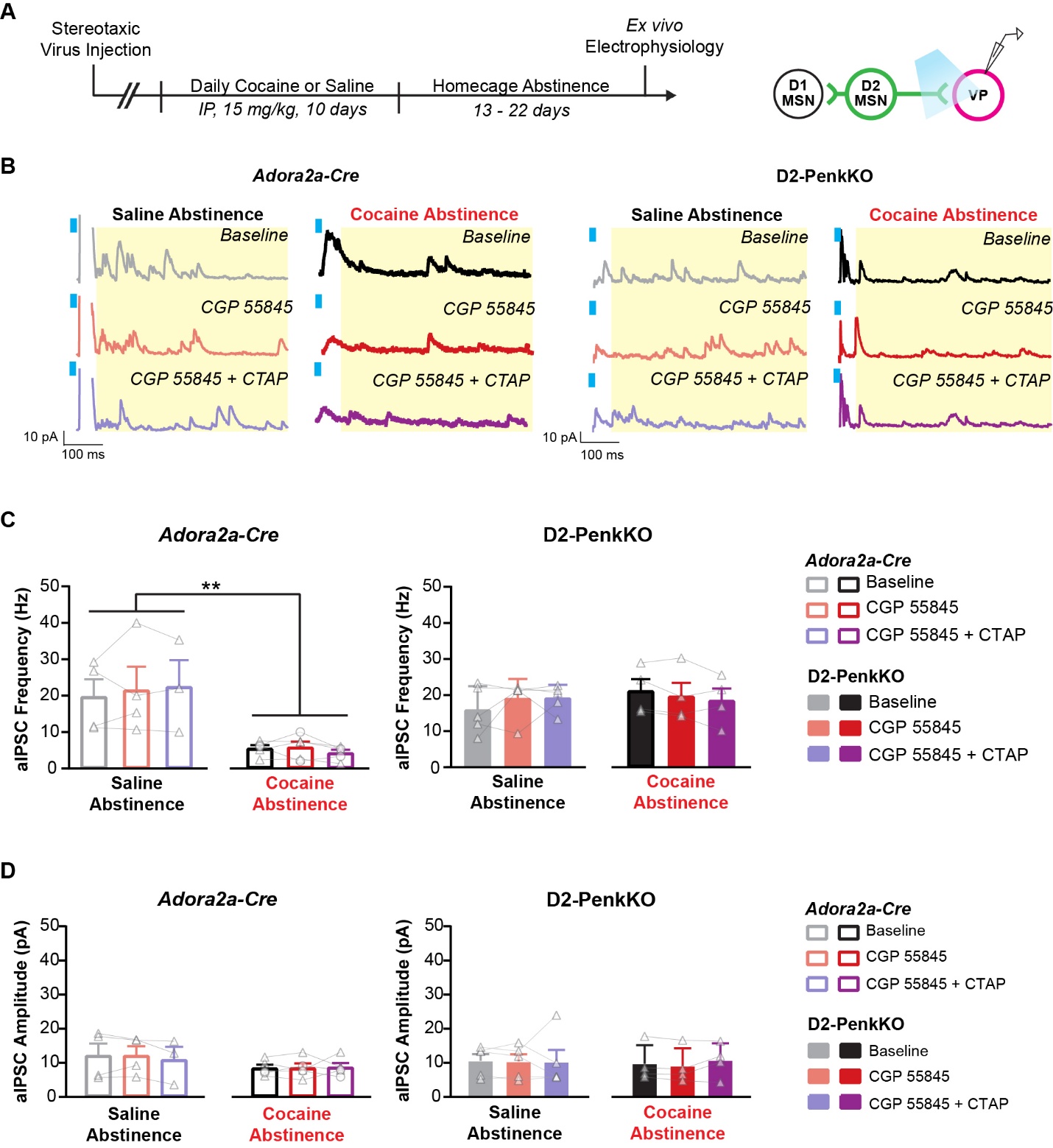

**Supplemental Figure 3. A GABA_B_ receptor antagonist does not reverse cocaine-abstinence-induced GABA suppression in the ventral pallidum.** (A) Experiment timeline (left) and electrophysiology recording configuration (right). (B) Representative aIPSC traces from saline- and cocaine-abstinent *Adora2a*-Cre controls (left) and D2-PenkKOs (right) at baseline and following bath application of the GABA_B_ antagonist CGP 55845 (2 µM, red) and CGP 55845 with the MOR antagonist CTAP (1 µM) (purple). The yellow box denotes the 50 – 500 ms analysis window after optogenetic stimulation (cyan). (C) Cocaine abstinence inhibited the average aIPSC frequency in *Adora2a*-Cre controls (left, 5 cells/2 mice) but not D2-PenkKOs (right, 4 cells/2 mice), relative to same-genotype saline-treated counterparts (saline D2-PenkKO: 5 cells/2 mice; saline *Adora2a*-Cre: 4 cells/2 mice). Neither CGP 55845 alone nor CGP 55845 + CTAP reversed cocaine-abstinence-induced GABA suppression. (D) Average aIPSC amplitude was not affected by cocaine abstinence, CGP 55845, or the combination of CGP 55845 with CTAP for either genotype. Data are shown as mean ± SEM, with individual data points labeled as male (circles) and female (triangles). ***p* < 0.01

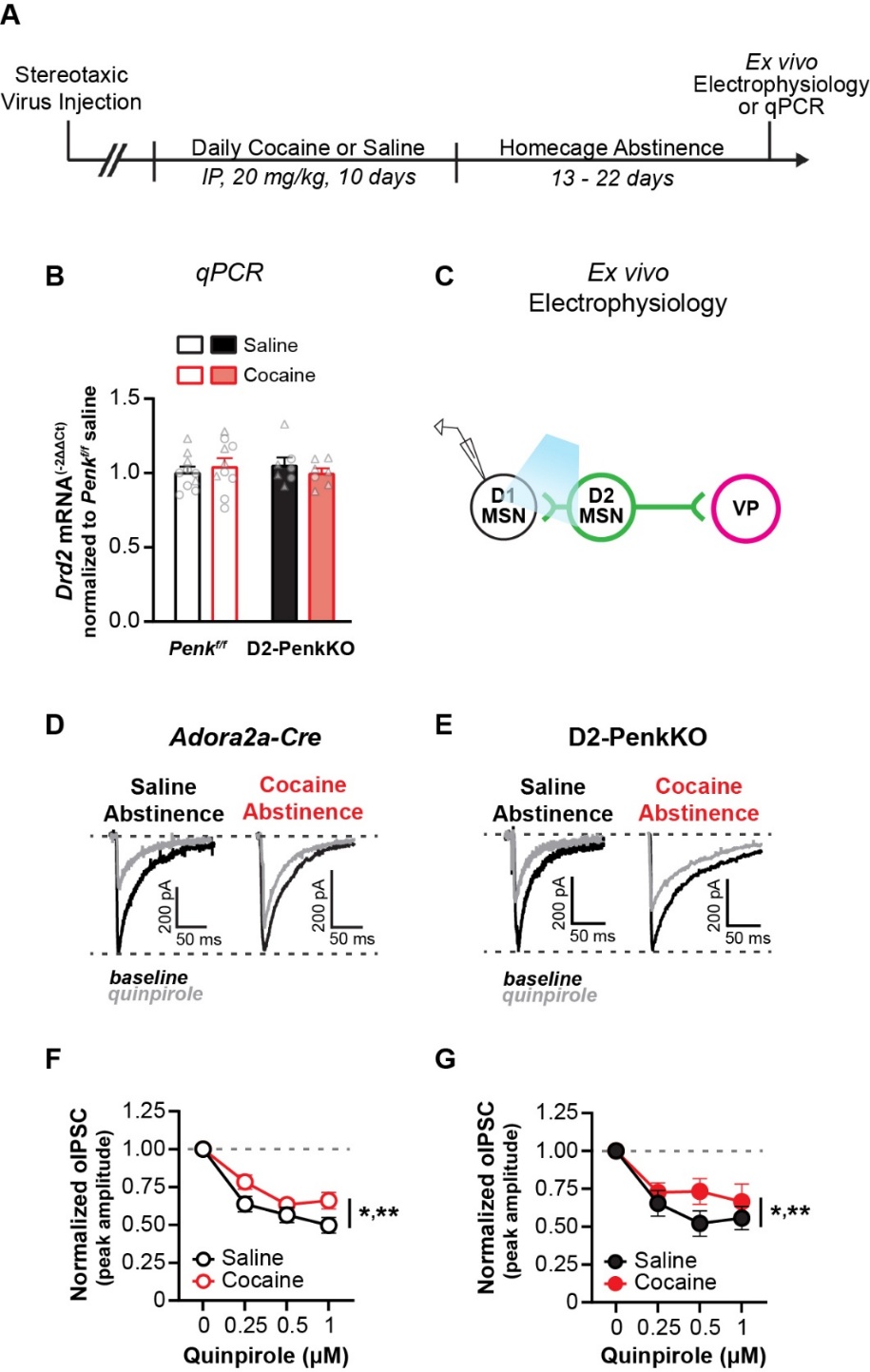

**Supplemental Figure 4. Cocaine abstinence, but not low striatal enkephalin, decreases the sensitivity of D2 receptors in D2-MSNs.** (A) Experiment timeline. (B) Cocaine abstinence does not alter mRNA abundance of the dopamine D2 receptor gene (*Drd2*) in D2-PenkKO mice (saline, n = 7; cocaine, n = 7) or littermate *Penk^f/f^* controls (saline, n = 10; cocaine, n = 10). (C) Electrophysiology recording configuration. (D, E) Example traces of optogenetically-evoked inhibitory post synaptic currents (oIPSCs) originating from D2-MSNs and recorded in D1-MSNs are shown for *Adora2a*-Cre controls (D) and D2-PenkKOs (E). Recordings were made from saline-abstinent (left) and cocaine-abstinent (right) subjects and are shown at baseline (black) and in the presence of quinpirole (gray). (F, G) Quinpirole dose-dependently inhibited oIPSC amplitude in *Adora2a*-Cre controls (saline: 18 cells/7 mice, cocaine: 13 cells/6 mice, F) and D2-PenkKOs (saline: 8 cells/3 mice, cocaine: 7 cells/3 mice) (Quinpirole x Cocaine, *p* < 0.0001, G). However, quinpirole was less effective at suppressing oIPSC amplitude in cocaine-abstinent mice compared to saline-abstinent mice. Asterisks represent post hoc t-tests comparing saline and cocaine groups collapsed across genotype. Saline vs. cocaine for 0.25 and 0.5 µM, * *p* < 0.05. Saline vs. cocaine for 1 µM, ** *p* < 0.01. Data are shown as mean ± SEM, with individual data points labeled as male (circles) and female (triangles).

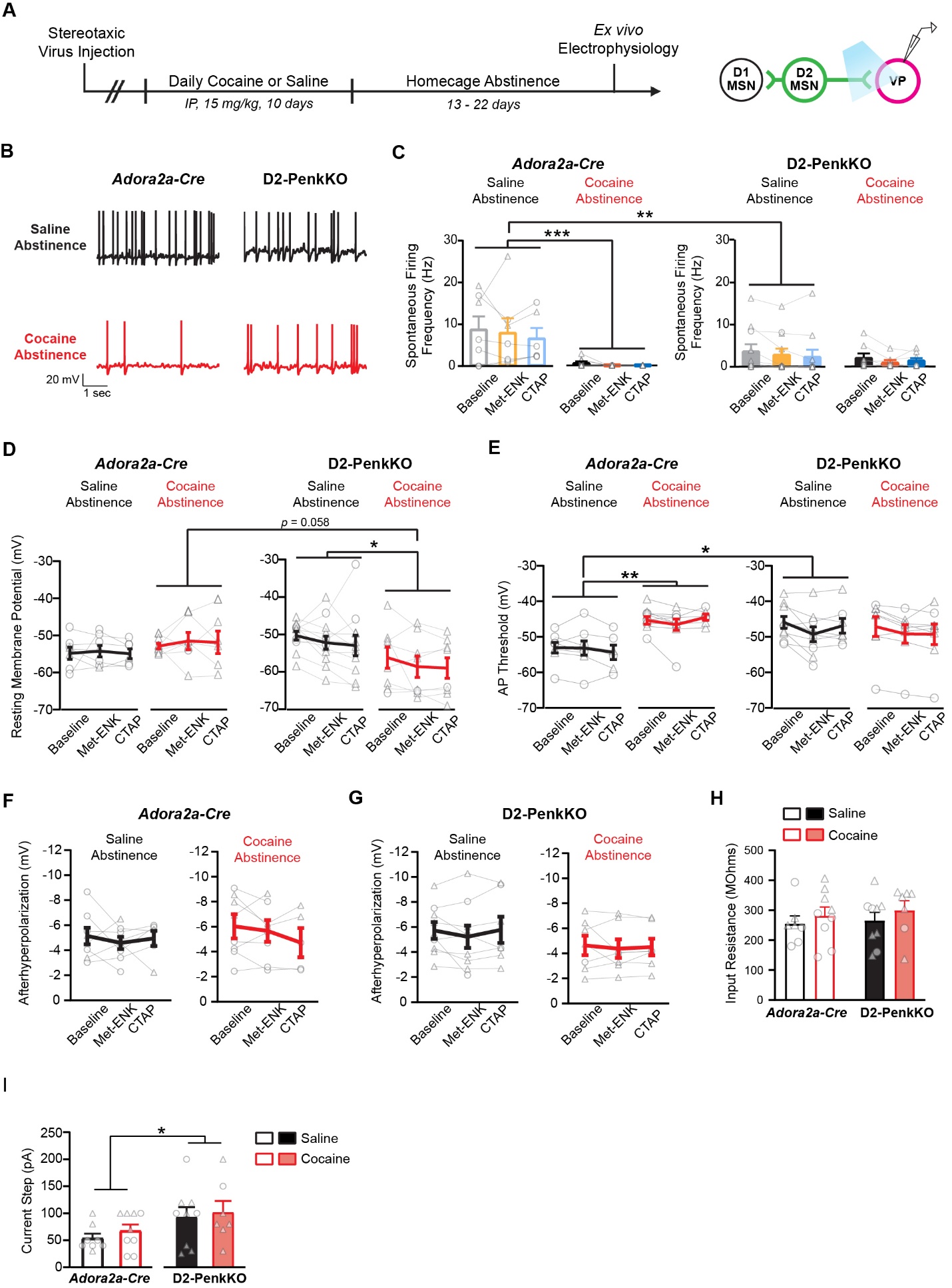

**Supplemental Figure 5. Cocaine abstinence and low striatal enkephalin induce adaptations in VP neuron excitability.** (A) Experiment timeline (left) and electrophysiology recording configuration (right). (B) Representative traces of baseline spontaneous firing of VP cells from saline-abstinent (black) or cocaine-abstinent (red) *Adora2a*-Cre controls (left) and D2-PenkKOs (right). (C) Spontaneous AP firing is reduced in cocaine-abstinent *Adora2a*-Cre controls (n = 9 cells/4 mice) and D2-PenkKOs (n = 8 cells/3 mice), as well as in saline-abstinent D2-PenkKOs (n = 11 cells/5 mice), compared to saline-abstinent *Adora2a*-Cre controls (n = 7 cells/3 mice). (D) Cocaine abstinence hyperpolarized the resting membrane potential in D2-PenkKOs compared to *Adora2a*-Cre controls and saline-abstinent D2-PenkKOs. (E) Action potential (AP) threshold was depolarized in cocaine-abstinent *Adora2a*-Cre controls (n = 9 cells/5 mice) and saline-abstinent D2-PenkKOs (n = 9 cells/5 mice) compared to saline-abstinent *Adora2a*-Cre controls (n = 8 cells/5 mice), but not cocaine-abstinent D2-PenkKOs (n = 7 cells/3 mice). (F-H) The after hyperpolarization of action potentials (F, G) and input resistance (H) of VP neurons are shown. There was no effect of met-enkephalin (Met-ENK, 1 µM), CTAP (1 µM), or genotype on either metric (saline *Adora2a*-Cre: 8 cells/5 mice; cocaine *Adora2a*-Cre: 8 cells/5 mice; saline- D2-PenkKO: 9 cells/5 mice; cocaine D2-PenkKO: 7 cells/3 mice). (I) D2-PenkKO VP neurons (saline and cocaine-abstinent) require more current injection to elicit ~ 10 action potentials compared to *Adora2a*-Cre controls (saline *Adora2a*-Cre: 8 cells/5 mice; cocaine *Adora2a*-Cre: 9 cells/5 mice; saline D2-PenkKO: 9 cells/5 mice; cocaine D2-PenkKO: 7 cells/3 mice). Data are shown as mean ± SEM, with individual data points labeled as male (circles) and female (triangles). * *p* < 0.05, ** *p* < 0.01, *** *p* < 0.001

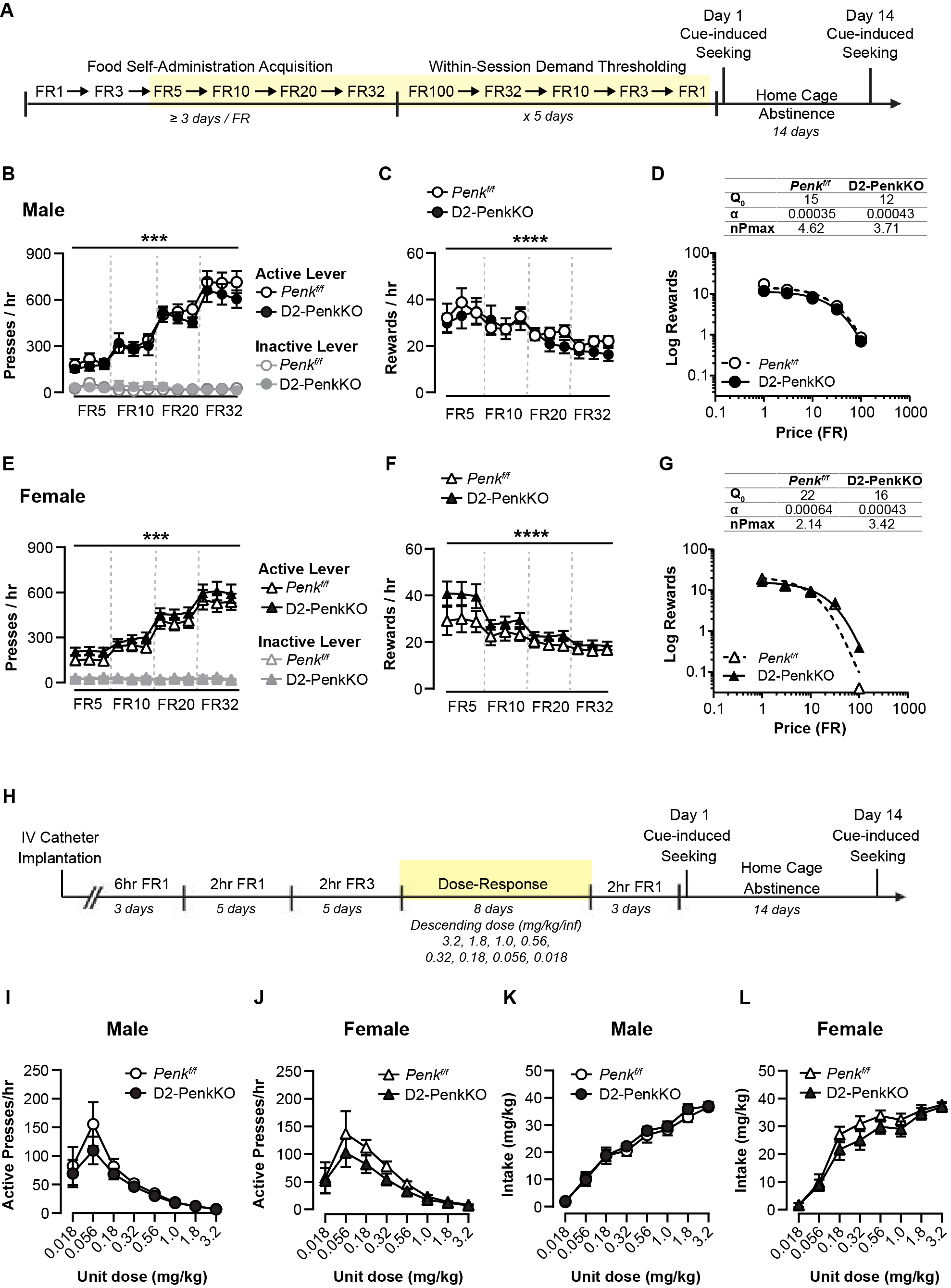

**Supplemental Figure 6. Deletion of striatal enkephalin does not affect self-administration of sucrose at high fixed ratios or cocaine self-administration across a dose response.** (A) Experimental timeline for sucrose self-administration. The yellow box denotes the experimental phases to which the displayed data correspond. (B, E) Active (black) and inactive (grey) lever presses per hour for sucrose pellets are shown for male (B, circles) and female (E, triangles) D2-PenkKO (closed symbols) and *Penk*^f/f^ littermate controls (open symbols). D2-PenkKO showed comparable responding and lever discrimination compared with controls over increasing fixed ratios (FR5, FR10, FR20, FR32). Average time to complete the entire FR progression was 21.1 ± 1.3 days. (C, F) Rewards earned per hour increased over trial similarly for males (C) and females (F) and between genotypes. (D, G) Demand curves generated based on the average of 5 days of a within-session thresholding procedure are shown. There were no differences between genotype or sex in the demand curve for earned sucrose pellets. There were also no differences in parameters (Q0, α, nPmax) estimated from these curves (inset tables). (H) Experiment timeline for cocaine self-administration. The yellow box denotes the experimental phases to which the displayed data correspond. (I, J) Active lever presses per hour across varying unit doses of cocaine are shown for males (I) and females (J). D2-PenkKOs (closed symbols) exhibited comparable responding compared to *Penk*^f/f^ controls (open symbols). (K, L) Total intake for each cocaine unit dose is shown for males (K) and females (L) and did not differ between genotypes. Data are shown as mean ± SEM, with group data points labeled as male (circles) or females (triangles). *** *p* < 0.001, **** *p* < 0.0001

**References**

1. Matsumura K, Nicot A, Choi IB, Asokan M, Le NN, Natividad LA, Dobbs LK (2023): Endogenous opioid system modulates conditioned cocaine reward in a sex‐dependent manner. *Addiction Biology* 28. https://doi.org/10.1111/ADB.13328

2. Remmers B, Nicot A, Matsumura K, Lyuboslavsky P, Choi IB, Ouyang Y, Dobbs LK (2025): Mu opioid receptors expressed in striatal D2 medium spiny neurons have divergent contributions to cocaine and morphine reward. *Neuroscience* 568: 273–284.

3. Schurmann B (2010): *Endogenous Opioid Peptides in Drug Addiction*. The University of Bonn.

4. Goldsmith JR, Perez-Chanona E, Yadav PN, Whistler J, Roth B, Jobin C (2013): Intestinal epithelial cell-derived μ-opioid signaling protects against ischemia reperfusion injury through PI3K signaling. *American Journal of Pathology* 182: 776–785.

5. Creed M, Ntamati NR, Chandra R, Lobo MK, Lüscher C (2016): Convergence of Reinforcing and Anhedonic Cocaine Effects in the Ventral Pallidum. *Neuron* 92: 214–226.

6. Mahler S V, Vazey EM, Beckley JT, Keistler CR, Mcglinchey EM, Kaufling J, *et al.* (2014): Designer receptors show role for ventral pallidum input to ventral tegmental area in cocaine seeking. *Nat Neurosci* 17: 577–585.

7. Dobbs LK, Kaplan AR, Lemos JC, Matsui A, Rubinstein M, Alvarez VA (2016): Dopamine Regulation of Lateral Inhibition between Striatal Neurons Gates the Stimulant Actions of Cocaine. *Neuron* 90: 1100–1113.

8. Dobbs LK, Kaplan AR, Bock R, Phamluong K, Shin JH, Bocarsly ME, *et al.* (2019): D1 receptor hypersensitivity in mice with low striatal D2 receptors facilitates select cocaine behaviors. *Neuropsychopharmacology* 44: 805–816.

9. Crespo JA, Manzanares J, Oliva JM, Corchero J, Palomo T, Ambrosio E (2001): Extinction of cocaine self-administration produces a differential time-related regulation of proenkephalin gene expression in rat brain. *Neuropsychopharmacology* 25: 185–194.

10. Mongi-Bragato B, Avalos MP, Guzmán AS, Bollati FA, Cancela LM, Toll L, *et al.* (2018): Enkephalin as a pivotal player in neuroadaptations related to psychostimulant addiction. *Front Psychiatry* 9: 1–11.

11. Dai KZ, Choi IB, Levitt R, Blegen MB, Kaplan AR, Matsui A, *et al.* (2022): Dopamine D2 receptors bidirectionally regulate striatal enkephalin expression: Implications for cocaine reward. *Cell Rep* 40: 111440.

12. Kircher DM, Aziz HC, Mangieri RA, Morrisett RA (2019): Ethanol experience enhances glutamatergic ventral hippocampal inputs to D1 receptor-expressing medium spiny neurons in the nucleus accumbens shell. *Journal of Neuroscience* 39: 2459–2469.

13. Bekkers JM, Clements JD (1999): Quantal amplitude and quantal variance of strontium-induced asynchronous EPSCs in rat dentate granule neurons. *Journal of Physiology* 516: 227–248.

14. Xu-Friedman MA, Regehr WG (2000): Probing fundamental aspects of synaptic transmission with strontium. *Journal of Neuroscience* 20: 4414–4422.

15. Gerdeman G, Lovinger DM (2001): CB1 cannabinoid receptor inhibits synaptic release of glutamate in rat dorsolateral striatum. *J Neurophysiol* 85: 468–471.

16. Hursh SR, Silberberg A (2008): Economic Demand and Essential Value. *Psychol Rev* 115: 186–198.

17. Bentzley BS, Fender KM, Aston-Jones G (n.d.): The behavioral economics of drug self-administration: A review and new analytical approach for within-session procedures. https://doi.org/10.1007/s00213-012-2899-2
